# Habitat and Hydrodynamics Influence Coral Reef and Seagrass Microbial and Exometabolite Dynamics

**DOI:** 10.64898/2026.01.12.699023

**Authors:** Brianna M. Garcia, Sharon L. Grim, Yan Jia, Laura Weber, Cynthia C. Becker, Mallory Kastner, Gretchen J. Swarr, Melissa C. Kido Soule, Aushaun Brown, Weifeng Zhang, Elizabeth B. Kujawinski, Amy Apprill

## Abstract

Coral reef and seagrass ecosystems provide critical storm protection and economic revenue to tropical coastal communities. Effective monitoring and restoration strategies are essential given increasing impacts from climate change and human development. Microorganisms, and the metabolites they produce and consume, are key drivers of coastal ecosystem function. However, microbially-mediated metabolite recycling remains poorly understood, limiting its inclusion in conservation and restoration strategies. Here we examine how seawater exometabolites and microorganisms in coastal ecosystems vary in relation to habitat (seagrass or coral reef), temporal scales and hydrodynamics. We characterized benthic seawater from two St. John, U.S. Virgin Islands coral reefs (Yawzi and Tektite) and one seagrass meadow at dawn and mid-day over four consecutive days. Using quantitative metabolomics and SSU rRNA gene amplicon sequencing, we found that habitat served as the primary driver of exometabolite and microbial community composition, while daily changes and hydrodynamics strongly influenced system variability. Hydrodynamic modeling indicated offshore water intrusion at mid-day at Yawzi reef, likely driving exometabolite and microbial shifts towards oligotrophic taxa (e.g., SAR11, SAR86). In contrast, a high percentage of coastal source water in the seagrass habitat maintained stable exometabolite pools and supported diverse microbial communities. These findings demonstrate that coastal habitat and hydrodynamics strongly influence exometabolite and microbial assemblages, with lesser contributions from temporal changes. Integrating exometabolites, microorganisms, and hydrodynamics provides new insights into coastal ecosystem functioning useful for habitat monitoring and restoration strategies.

## Introduction

Tropical coastal ecosystems, including coral reefs and seagrass habitats, support the livelihoods of around a billion people worldwide through services such as storm protection, sediment stabilization, tourism, and carbon sequestration [1, 2]. These ecosystems are under significant and rapidly increasing threat due to climate change and human development [3–5]. Understanding and monitoring the health of coral reefs and seagrass habitats is important for informing conservation and remediation activities and for analyzing future risks facing coastal regions. Current monitoring programs predominantly use macroorganismal metrics, such as the diversity and abundance of stony corals, algae, and fish as sentinels of ecosystem health [6]. However, visual observations miss critical information about the microbial and chemical parameters that play into ecosystem health in a dynamic ocean [6, 7].

Marine microbes are primary producers and consumers of dissolved organic matter (DOM), form symbiotic and/or pathogenic relationships with corals and other macroorganisms, and have the potential for rapid growth and cellular turnover [8]. The most dynamic component of DOM, the pool of readily consumed molecules, is composed of structurally diverse, low molecular weight molecules (exometabolites) that are rapidly transformed through microbial metabolism [8]. This tightly interconnected microbe-metabolite network can reflect real-time changes in environmental conditions, including nutrient and pollution levels, oxygen, temperature, and salinity [9–12]. Additionally, physical processes such as hydrodynamics can influence exometabolite and microbial dynamics within tropical coastal ecosystems. For example, reefs situated along the forereef are more exposed to offshore seawater, and likely experience different inputs and turnover rates of exometabolites compared to reefs located in more protected bays. Understanding the influence of hydrodynamics on differing metabolite inputs can shed light on the tightly coupled responses of microbial growth and succession in seawater. Despite the importance of examining interactions between seawater exometabolites, microorganisms, and hydrodynamics, no studies have yet examined these three parameters simultaneously in tropical coastal environments.

Over the last decade, investigations have established a taxonomic knowledge of microorganisms within tropical coastal ecosystems. Seawater microorganisms on shallow coral reefs include high standing stocks of photoautotrophic cells (e.g., *Prochlorococcus*, *Synechococcus*) and varying densities of heterotrophic bacterial taxa (oligotrophs: e.g., SAR11, SAR86, NOR5; and copiotrophs: e.g., *Flavobacteriaceae*, *Rhodobacteraceae*) [13]. Reef site and biogeography serve as the primary control on microbial composition [14–18], likely due to the combination of ocean and reef conditions. Additional controls include macro-organismal density and diversity [19–22], physicochemical conditions of the seawater [23–25], and disturbance events such as disease [26, 27]. For seagrasses, microbial composition is linked to site as well as tidal conditions and salinity [28–30]. The largest within-site impact on temporal dynamics within reefs and seagrass environments appears to be the nightly division of *Prochlorococcus* and *Synechococcus* cells [28, 31, 32]. Collectively, seawater microorganisms can serve as bioindicators of tropical coastal ecosystems and ocean conditions, and there is momentum to provide a more unified platform for monitoring programs [7, 33].

Compared to seawater microorganisms, much less is known about the exometabolome composition and dynamics in coral reef and seagrass habitats. Metabolomics studies have largely focused on the intracellular composition of coral tissues and seagrass leaves, roots, and rhizomes; relatively few studies have addressed the chemical composition and spatio-temporal dynamics of their exometabolomes [16, 34–36]. Like microbial communities, DOM composition varies with biogeography [16], disturbance [37], and anthropogenic and terrestrial influences [17, 38]. Coral reef primary producers release distinct chemicals with diurnal impacts on microbial communities [20] and water chemistry parameters, such as dissolved oxygen, pH, and nutrient availability [39]. Specific chemical classes differ on diel timescales based on the benthic producer, with coral and fleshy algae found to release more bioavailable lipids and organic nitrogen compounds at night and labile heterocyclic compounds during day [40]. Research on seagrass exometabolomes is more limited, however studies indicate that eelgrass leaf exudates can contribute substantially to marine DOM, particularly with allelochemicals [35]. Quantitative metabolomic studies of coral reefs and seagrass meadows have been limited by methodological constraints. However, emerging approaches such as chemical derivatization support quantification of new exometabolites as well as the flux dynamics of individual compounds or compound classes [34, 41, 42].

Translating molecular-level insights into ecosystem-scale processes requires consideration of hydrodynamics and physical geography, because water flow strongly influences microbial activity and metabolite transfer [43–45]. Coastal circulation directly affects nutrient delivery, disease transport, contaminant distribution, and larval dispersal [45]. Numerical simulations suggest a pattern of increased nutrient uptake near the reef crest, with decreasing nutrient concentrations downstream across the reef [43], underscoring that local flow patterns shape resource availability. Empirical studies support these patterns. For example, dissolved organic carbon (DOC) and microbial biomass are depleted on fringing and barrier reef habitats in Mo’orea, French Polynesia, alongside differentiation of bacterioplankton communities between offshore, forereef, backreef, and bay environments [46]. Similarly, Weber et al. observed high biogeochemical similarity across forereefs in the Jardines de la Reina, Cuba reef system, hypothesizing that regional-scale hydrodynamics, such as the Caribbean current, may homogenize exometabolite pools by flushing reefs with oligotrophic water [36]. Together, these examples illustrate that water flow can either concentrate or dilute microbial–metabolite interactions, yet the extent to which hydrodynamics mediate ecosystem-scale chemical diversity and microbial community composition remains poorly resolved. Addressing this gap requires systems where microbial, exometabolite, and hydrodynamic measurements can be directly integrated.

The coral reefs of southern St. John, US Virgin Islands, present an ideal framework to examine exometabolite-microbial-hydrodynamic patterns. Extensive studies over four decades provide a rich ecological context for understanding current community dynamics relative to well-documented coral loss and habitat change [47–50]. The microorganisms on these reefs and adjacent seagrass meadows have been studied for 10 years [18, 26, 28, 31] and the exometabolites recently explored [20, 34], establishing a baseline of microbial and chemical diversity. Further, a realistic high-resolution hydrodynamic model of the St. John coastal region was recently constructed and validated, and used to calculate the cross-bay variability of particle residence time, a key factor influencing nutrient delivery, disease spread, larval dispersal, and contaminant transport [44, 45]. Building on this extensive historical and environmental knowledge, this study examined the composition and diversity of seawater microorganisms and exometabolites in a well-characterized tropical embayment in relation to relevant influences, including site, habitat type, time of day, and particle origins. Over four days, we collected benthic depth seawater at dawn and mid-day at two coral reefs and one seagrass habitat. Seawater was processed and analyzed for dissolved metabolites using a benzoyl chloride (BC) derivatization approach followed by quantitative metabolomics [41], and for microbial community composition using small subunit (SSU) ribosomal RNA gene amplicon sequencing targeted to bacteria and archaea. Additionally, a hydrodynamic model previously established for this region and an associated particle-tracking model were used to trace back the sources of the water samples over the four sampling days. Our results reveal that habitat type (coral reef vs. seagrass), hydrodynamics, and time of day (dawn vs. mid-day), all influence the observed microbial-metabolite patterns.

## Materials and Methods

### Sample collection

Seawater samples were collected in January 2021 in St. John, U.S. Virgin Islands within the Virgin Islands National Park (Figure 1) at two coral reefs, Tektite (n=25, 18.309°N, 64.723°W, 8.4 m average collection depth) and Yawzi (n=25, 18.314°N, 64.726°W, 8.5 m), and one seagrass meadow (n=24, 18.317°N, 64.723°W, 3.6 m). Sampling locations were selected based on their proximity within the Great Lameshur Bay region and their unique benthic compositions determined via point intercept benthic surveys [34]. Samples were collected at ∼6 am (dawn) and ∼2 pm (mid-day) local time over four consecutive days. Scuba divers collected benthic seawater (within 0.25 m of the benthos) in acid-washed Niskin Bottles (General Oceanics) at three or four distinct locations at each site. After ascent, the Niskins were drained into individual polycarbonate (PC) bottles, placed on ice, and processed within 2 h of collection. Each individual Niskin bottle was considered a representative sample of each site (biological replicate, herein defined as a replicate), with a total of 3-4 replicates collected per site at each sampling timepoint.

**Figure 1.**
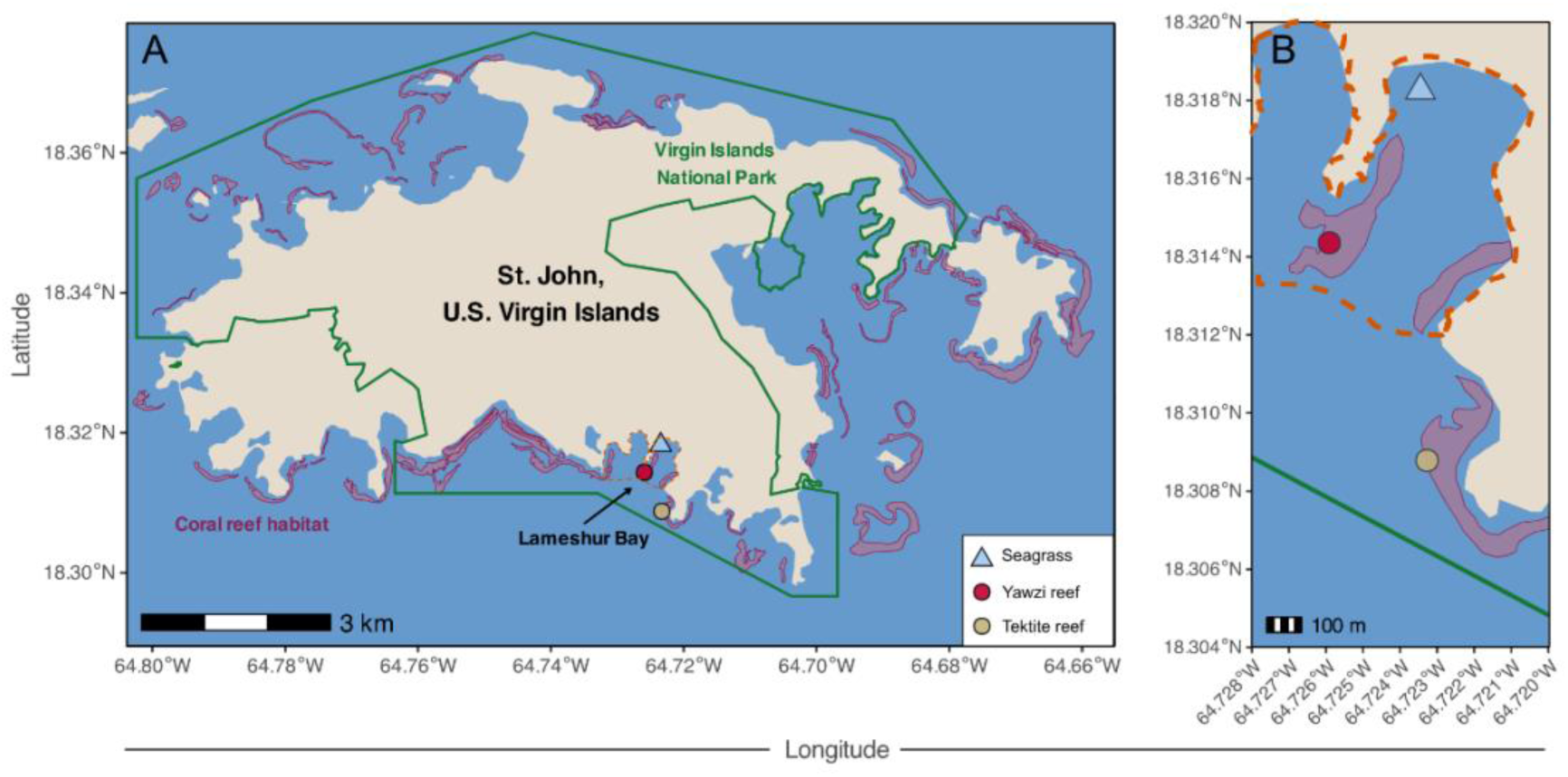
Map of St. John, U.S. Virgin Islands. (tan) showing (A) the Virgin Islands National Park (green boundary) and surrounding coral reefs (pink). Sampling sites are indicated for two coral reef habitats, Yawzi reef (red circle) and Tektite reef (brown circle), and one seagrass meadow (blue triangle). The boundary of the larger Lameshur Bay region is shown (orange, dashed line). (B) Enlarged view of sampling sites within the Great Lameshur Bay region. Map shapefiles were obtained from open-access sources: Stanford EarthWorks (island boundary), the National Park Service Land Resources Division (park boundary), and NOAA National Centers for Coastal Ocean Science (benthic habitat map).

### Filtration for microbial and metabolomics sample processing

Benthic seawater was filtered through polytetrafluoroethylene 0.2-μm, 47-mm filters (Omnipore) using peristalsis (MasterFlex L/S pump and pump heads). Thermoplastic elastomer tubing and acid-washed fluorinated ethylene propylene tubing (890 Tubing, Nalgene) were used to pump seawater through the filter membrane and into acid-washed PC collection bottles. Filters with captured cells were placed in cryovials and flash frozen in a dry shipper, where they remained stored until transport to Woods Hole Oceanographic Institution (WHOI) and storage at -80°C until processing. Corresponding filtrate (40 mL) was collected into combusted amber vials for BC derivatization and targeted metabolomics measurements. Metabolomics samples were stored and transported at -20°C.

### Exometabolite sampling processing

Filtered, frozen seawater samples were thawed and 25 mL from each sample was transferred into a combusted 40 mL amber glass vial for ^12^C-benzoyl chloride (^12^C-BC) derivatization conducted according to Widner et al. 2021 [41]. A nine-point standard curve was derivatized (^12^C-BC) in parallel with samples, using 25 mL of off-reef seawater as the matrix and included the values: 0, 0.125, 0.25, 0.75, 1.75, 2.5, 7.5, 17.5, and 25 ng-added. Stable isotopically labeled internal standards (^13^C-labeled SIL-IS) were derivatized in parallel, using ^13^C-BC, and spiked into all experimental and standard curve samples. Derivatization methods are described in detail elsewhere [34]. Derivatized samples and standards were then extracted by SPE using 6 mL, 1 g Bond Elut PPL cartridges (Agilent), and the final eluent was dried via vacufuge to near-dryness at 30°C. Dried samples were stored at -20°C until mass spectrometry analysis, at which time each sample was reconstituted in 100 µL of 5% acetonitrile (MeCN), transferred to a 2 mL vial with a small volume insert, and topped with 5 μL of 100% MeCN. Samples were stored at 4°C until instrumental analysis.

### Microbiome sample processing

DNA was extracted from seawater filters using DNeasy PowerBiofilm Kits (Qiagen) according to the manufacturer’s protocol, with a blank (control) per extraction round. The eluted DNA served as the template for barcoded PCR reactions to amplify the hypervariable 4 region (V4) of the 16S SSU rRNA gene using the 515F [51] and 806R [52] primers, respectively. The 50μL reactions contained: 2 μL DNA template (1:10 diluted), 0.5 μL of GoTaq DNA Polymerase, 1 μL each of forward and reverse primers at 10 μM, 1 μL of 10 mM deoxynucleoside triphosphate (dNTP) mix, 5 μL MgCl_2_, 10 μL GoTaq 5X colorless flexi buffer (all reagents Promega), and 29.5 μL nuclease-free water. Genomic DNA from Microbial Mock Community B (Even, Low Concentration), v5.1L, for 16S rRNA Gene Sequencing, HM-782D, was used as our sequencing control. Nuclease-free PCR-grade water was our negative PCR controls. PCR reactions were as follows: 95°C for 2 min; 29, 30, 32, or 34 cycles of 95°C for 20 s; 55°C for 15 s; 72°C for 5 min; and finally, 72°C for 10 min, before a hold at 4°C; cycle number varied to optimize amplification. DNA extraction kit controls were amplified at the highest PCR cycle number of any of the samples in each extraction batch. The barcoded PCR products were purified using the MinElute PCR Purification kit (Qiagen) and their concentrations were measured using Qubit 2.0 fluorometry. Purified PCR products were diluted to 1 ng/μL and pooled. Samples were sequenced 2×250bp on an Illumina MiSeq at the Roy J. Carver Biotechnology Center of the University of Illinois at Urbana-Champaign.

### Exometabolite and microbial data analysis

Metabolomic and 16S rRNA gene amplicon data were processed using established workflows in Skyline, MATLAB®, and R. Exometabolome and microbiome profiles were compared using Bray–Curtis and Morisita–Horn dissimilarities, respectively, visualized by non-metric multidimensional scaling (NMDS), and tested via PERMANOVA. Beta-diversity and beta-dispersion analyses quantified within- and between-site variability, with Kruskal–Wallis and pairwise Wilcoxon rank-sum tests assessing group differences. Comparisons were conducted based on site (Tektite vs. Yawzi vs. seagrass), habitat-type (seagrass vs. reef), sampling time (dawn vs. mid-day), and sampling day. Spearman correlations identified relationships between microbial taxa relative abundances and exometabolite concentrations. Linear regressions were conducted between hydrodynamic simulation outputs and exometabolome and microbiome variability using beta-dispersion distances. Detailed methods are available in the Supplemental Text.

### Hydrodynamic modeling

We used an offline particle tracking model, ROMSPath [53], to identify the sources of the water samples. Three-dimensional hourly velocity field from the hydrodynamic model 2021 simulation (see Supplemental Text) was used in ROMSPath to calculate the backward particle trajectories with a time step of 60 s. A total of 24 particle-tracking backward simulations were carried out per water sample. In each simulation, particles were released every 10 minutes for a two-hour window centered at a sampling time and then tracked for 24 hours to capture day-to-day and diurnal influences on particle trajectories. The released particles were evenly spaced every 2 m horizontally across a 50 m × 40 m area centered on each sampling station and every 1 m vertically over the whole water column. The numbers of particles within the larger Lameshur Bay region at each time was counted. The larger Lameshur Bay region was selected to represent a microbial environment distinct from the open coastal ocean. This boundary was chosen to focus our study on sites under stronger coastal influence, which are known to harbor characteristic microbial communities. Additionally, defining this region constrains our analysis to the Virgin Islands National Park, excluding reef sites outside this boundary that have previously been shown to possess unique exometabolome and microbiome profiles [28, 34]. Because of the high spatial density of released particles, the percentage of particles originating from the larger Lameshur Bay region at each time point is considered representative of the proportion of water samples derived from this area.

## Results

### Coral reefs and seagrass meadows harbor distinct pools of polar exometabolites and microbial communities

We quantified 45 amine- and alcohol-containing exometabolites with concentrations spanning roughly four orders of magnitude, ranging from 5 picomolar (pM) to 30 nanomolar (nM; Table S1). All 45 exometabolites were quantified in the seagrass meadow, whereas 42 were quantified at both reefs (Figure 2). For the seawater microbial community, an average of 87,355 sequences (range of 13,942 – 118,596; similar sequencing effort between sites and sampling times, p = 0.43 via Kruskal-Wallis) and 169-843 amplicon sequence variants (ASVs) per sample were recovered (Table S2). Of the distinctive taxa identified, 30 ASVs were responsible for at least 82.4% of the sequences per sample (Figure 2, Table S3), representing consistency across samples.

**Figure 2.**
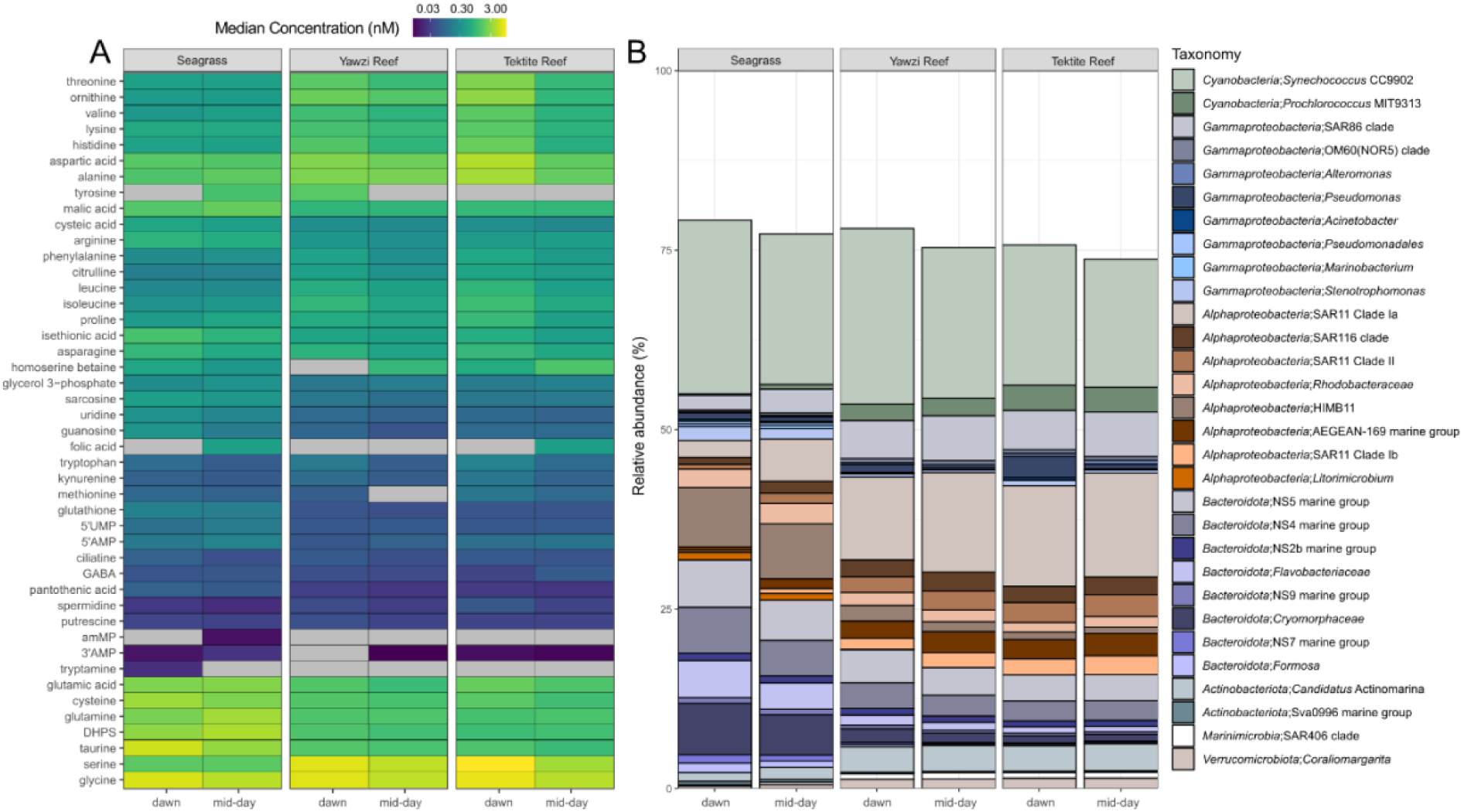
Exometabolites and microbes in coastal seawater sites. A) Median concentrations (nM) of quantified exometabolites (y-axis) in seagrass, Yawzi reef, and Tektite reef (n = 3-4) within a given sampling time (x-axis) across the four-day sampling period. Median concentrations are shown using a log-scaled color gradient to facilitate visualization across the wide dynamic range. Concentrations that fell below the limit of detection are represented by grey-filled boxes. Exometabolites were arranged along the y-axis according to hierarchical clustering of concentration values. B) Taxonomy bar chart of the 30 dominant lineages (as specific as genus) observed in seagrass, Yawzi reef, and Tektite reef. Each bar represents the sum of replicates (n = 2-3) for each sampling time, then relative abundance of the relevant taxa was calculated.

An unsupervised NMDS approach showed that exometabolite and microbial profiles were habitat-specific with reef sites clustered together and separate from the seagrass, irrespective of sampling time (Figure 3a,b). While clustering patterns by NMDS were similar for the exometabolome and microbiome, the samples’ beta diversity distributions (dispersion) were visibly and quantitatively different within the two ordination plots (Figure 3c,d). A PERMANOVA analysis identified the significant factors (p < 0.05) driving the exometabolome and microbiome variability observed in the NMDS plot (Table 1); site was the largest significant factor structuring both the exometabolome and microbiome, explaining 31.3% and 47.3% of the variability, respectively. Exometabolome and microbiome profiles of Tektite and Yawzi reefs had significantly different diversity distributions (bootstrapped pairwise t-tests, exometabolome t = 4.97, microbiome t = 2.93, p < 0.01), and the seagrass site was significantly different from each reef site (exometabolome t = -7.97 to -13.9, microbiome t = 6.50 to 9.40, p < 0.001, Table S4).

**Figure 3.**
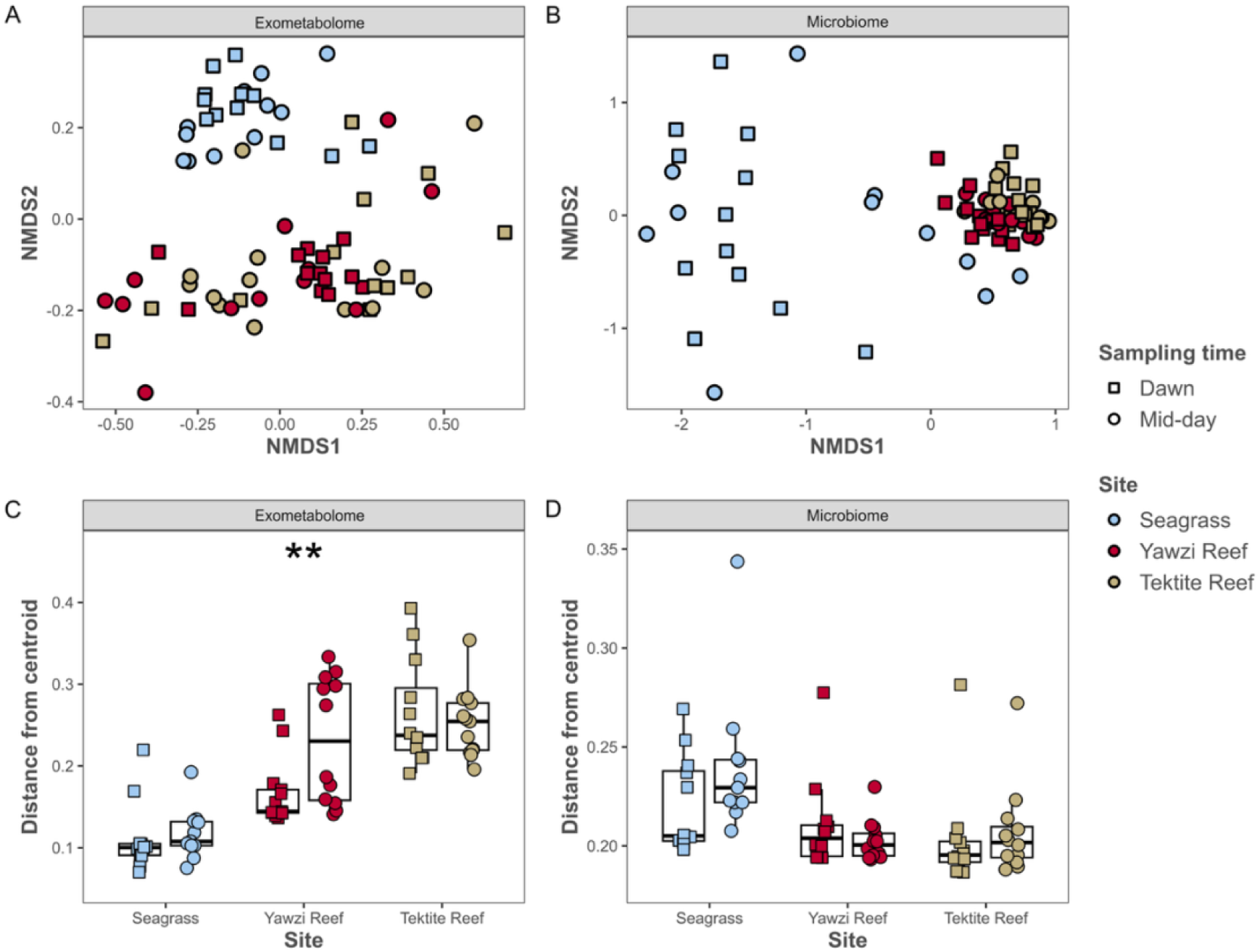
Coral reefs and seagrass harbor distinctive exometabolomes and microbiomes. NMDS of microbiomes from seagrass (blue), Yawzi reef (red), and Tektite reef (yellow). (A) Bray-Curtis dissimilarities between samples’ exometabolite concentrations were used to generate the NMDS. (B) Morisita-horn dissimilarities between samples’ ASV relative abundance profiles were used to generate the NMDS. PERMANOVA results are presented in Table 1. (C) Box-and-whisker plot showing beta-dispersion distances for the exometabolome data within each site and sampling time. Each point represents the difference in Bray-Curtis dissimilarity of a sample from the median dissimilarity (50th percentile line) for the time- and site-specific group. Intra-site and inter-site significant differences in dissimilarity distances are indicated with “**” (pairwise t-test p < 0.05). (D) Box-and-whisker plot showing beta-dispersion distances for the 16s rRNA sequencing data within each site and sampling time. Each point represents a sample’s difference in Morisita-Horn dissimilarity from the median dissimilarity (50th percentile line) for the time- and site-specific group.

**Table 1.** PERMANOVA and RDA results for exometabolome and microbiome.

|  | Exometabolomes |  |  |  | Microbiomes |  |  |  |
| --- | --- | --- | --- | --- | --- | --- | --- | --- |
| Source | Df | Pseudo-F | P-value | % of variation explained in model | Df | Pseudo-F | P-value | % of variation explained in model |
| Site | 2 | 15.6 | **0.0001** | 31.3 | 2 | 78.3 | **0.0001** | 47.3 |
| Time | 1 | 3.98 | **0.016** | 15.8 | 1 | 15.7 | **0.0002** | 21.2 |
| Day | 3 | 1.31 | 0.246 | 9.1 | 3 | 7.59 | **0.0007** | 14.7 |
| Day x Site | 6 | 2.17 | **0.011** | 11.7 | 6 | 2.42 | **0.039** | 8.3 |
| Time x Day | 3 | 1.25 | 0.270 | 8.9 | 11 | 0.342 | 0.95 | 3.1 |
| Time x Site | 2 | 1.11 | 0.353 | 8.4 |  |  |  |  |
| Time x Day x Site | 6 | 0.753 | 0.728 | 6.9 |  |  |  |  |
| Residual | 49 |  |  | 7.9 | 47 |  |  | 5.3 |
| Total | 72 |  |  |  | 70 |  |  |  |

Of the 45 quantified exometabolites, 34 showed significantly differentially abundant (SDA) concentrations between sites, within each sampling time. Specifically, 32 metabolites differed significantly at dawn and 29 at mid-day (false discovery rate [FDR] -adjusted P < 0.05, Pairwise Wilcoxon Rank Sum Test; Table S6). Exometabolite concentrations were largely elevated in the seagrass site, with only 11 metabolite concentrations significantly increased at either Yawzi or Tektite reef relative to the seagrass. The seagrass exometabolome was generally characterized by elevated concentrations of nucleotides (e.g., 3’AMP, 5’UMP), organosulfur compounds (e.g., DHPS, taurine), and stress-protective osmolytes (e.g., homoserine betaine). In contrast, coral reef exometabolomes were dominated by proteinogenic amino acids such as isoleucine, leucine, phenylalanine, serine, and threonine. Pair-wise exometabolite differences between the two reef sites were subtle, with only three SDA exometabolites. We conclude that the observed site differences were largely attributed to habitat type rather than individual site. A similar habitat-driven pattern was evident in the microbiome. Across reef and seagrass sites, 181 ASVs differed significantly in relative abundance based on concordance between corncob (5% FDR, p < 0.054) and DESeq2 (10% FDR, p < 0.050, Table S7, Table 2). Reef sites were enriched in cyanobacteria (i.e., *Prochlorococcus*, *Synechococcus*), oligotrophic heterotrophs such as *Candidatus* Actinomarina, and SAR11. Seagrass sites, in contrast, showed higher abundances of heterotrophic bacteria including *Rhodobacteraceae*, multiple Bacteroidia lineages (NS4, NS5, NS9), and *Cryomorphaceae*. These patterns aligned with cell counts, which showed that *Synechococcus* populations were 2-3x more abundant than *Prochlorococcus* overall, and consistently elevated in reef sites relative to seagrass (Figure S1b, Table S8). Like the exometabolites, the sequencing data pointed to habitat as the main distinguishing driver, clearly distinguishing reefs from seagrass while showing subtle differences among reefs (Figure 3).

**Table 2.**
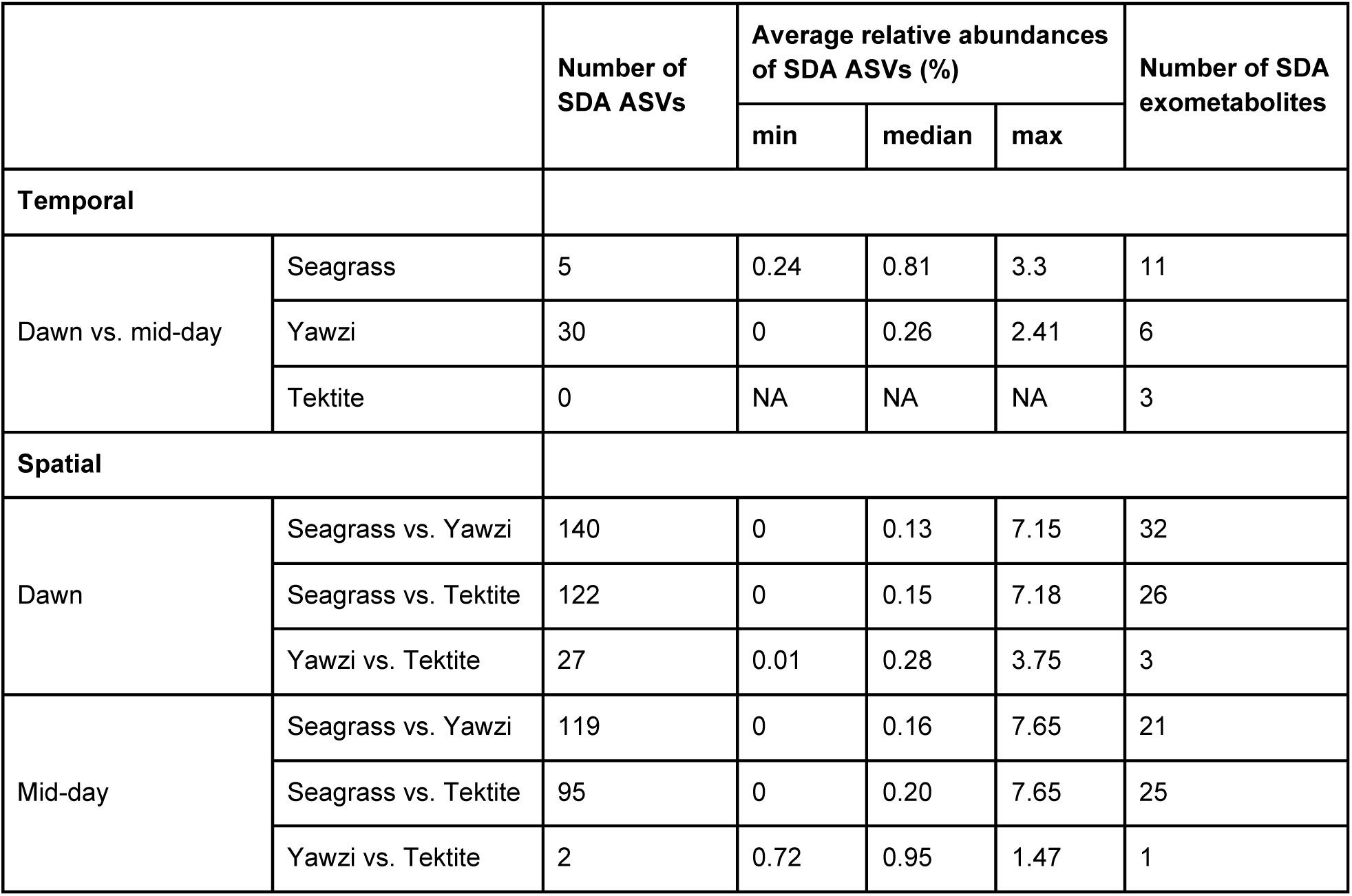
Frequency of significantly differentially abundant (SDA) ASVs and exometabolites across comparisons.

|  |  | Number of SDA ASVs | Average relative abundances of SDA ASVs (%) |  |  | Number of SDA exometabolites |
| --- | --- | --- | --- | --- | --- | --- |
|  |  |  | min | median | max |  |
| Temporal |  |  |  |  |  |  |
| Dawn vs. mid-day | Seagrass | 5 | 0.24 | 0.81 | 3.3 | 11 |
|  | Yawzi | 30 | 0 | 0.26 | 2.41 | 6 |
|  | Tektite | 0 | NA | NA | NA | 3 |
| Spatial |  |  |  |  |  |  |
| Dawn | Seagrass vs. Yawzi | 140 | 0 | 0.13 | 7.15 | 32 |
|  | Seagrass vs. Tektite | 122 | 0 | 0.15 | 7.18 | 26 |
|  | Yawzi vs. Tektite | 27 | 0.01 | 0.28 | 3.75 | 3 |
| Mid-day | Seagrass vs. Yawzi | 119 | 0 | 0.16 | 7.65 | 21 |
|  | Seagrass vs. Tektite | 95 | 0 | 0.20 | 7.65 | 25 |
|  | Yawzi vs. Tektite | 2 | 0.72 | 0.95 | 1.47 | 1 |

### Fine-scale temporal dynamics of the microbiome and exometabolome

While habitat was the main driver of community differences in both the exometabolome and microbiome, sampling time was also a large contributor, explaining more than 15% of the observed variability (Table 1). Site-specific NMDS ordinations with 95% confidence intervals showed little separation by time for either the exometabolome or microbiome (Figure S2a,b), with the largest temporal separation observed in the seagrass exometabolome, although confidence intervals of the two groups overlapped.

Diel changes in the exometabolomes and microbiomes showed differing dissimilarity patterns (Table S4). We calculated the mean difference in beta diversity, here reported as Δ*β*, from groups of sample beta diversity values to quantify whether communities became more or less dissimilar between dawn and mid-day. In the seagrass, microbial diversity significantly increased at mid-day (Δ*β* = +0.064, t = -4.27, p < 0.001), while exometabolome diversity did not change (Δ*β* = +0.005). No significant temporal shifts were detected in either microbiome (Δ*β* = 0.015) or exometabolome diversity (Δ*β* = -0.028) at Tektite reef. At Yawzi reef, microbial diversity was significantly lower at mid-day (Δ*β* = -0.031, t = 3.09, p = 0.003), while exometabolome diversity was significantly higher at mid-day (Δ*β* = +0.125, t = -6.72, p < 0.001), highlighting a unique pattern of opposing dynamics between microbial and exometabolite communities at this site.

Beta-diversity dispersion was used to quantify the spread of samples within each time point (Figure 3c,d, Table S5), providing a measure of how variable or homogeneous samples were relative to each group’s centroid. In the seagrass-associated communities, there was less dispersion in the exometabolome samples and relatively more dispersion in the corresponding microbiomes. The opposite pattern emerged for the coral reefs (Figure 3c,d). For the exometabolome, beta-dispersion (here represented as *σ*_*β*_^2^) increased from dawn to mid-day and was roughly two-fold higher in the reef sites compared to the seagrass. While exometabolomes across all sites, as well as the microbiomes of Tektite reef and the seagrass site, showed this general increase in heterogeneity from dawn to mid-day, the increase was only significant at Yawzi reef (p < 0.05; median dawn *σ*_*β*_^2^ = 0.144; median mid-day *σ*_*β*_^2^ = 0.230) (Figure 3c). In contrast, Yawzi reef microbiome beta dispersion values were generally lower at mid-day compared to dawn, although not significantly (median dawn *σ*_*β*_^2^ = 0.204; median mid-day *σ*_*β*_^2^ = 0.200) (Figure 3d).

Diel variation in exometabolite concentrations and microbial ASV relative abundances was observed across the three sites (Table 2); however, patterns varied in magnitude and direction (Figure 4). Across all sites, 15 exometabolites and 33 ASVs showed significant temporal trends (Table S6-7). The seagrass exometabolome had the largest number of SDA exometabolites (11) followed by Tektite (6), and Yawzi reef (3). Among the 15 significantly temporal exometabolites, cysteine and homoserine betaine were the only exometabolites found to exhibit significantly diel trends at all three sites. Additionally, homoserine betaine was the only exometabolite to show differing trends between sites (i.e., homoserine betaine was significantly elevated at dawn in the seagrass but was elevated at mid-day at both Tektite and Yawzi reefs). In contrast, cysteine concentrations were consistently increased at dawn compared to mid-day across all three sites, however, concentrations in the seagrass at both dawn and mid-day were significantly higher in the seagrass than either coral reef site.

**Figure 4.**
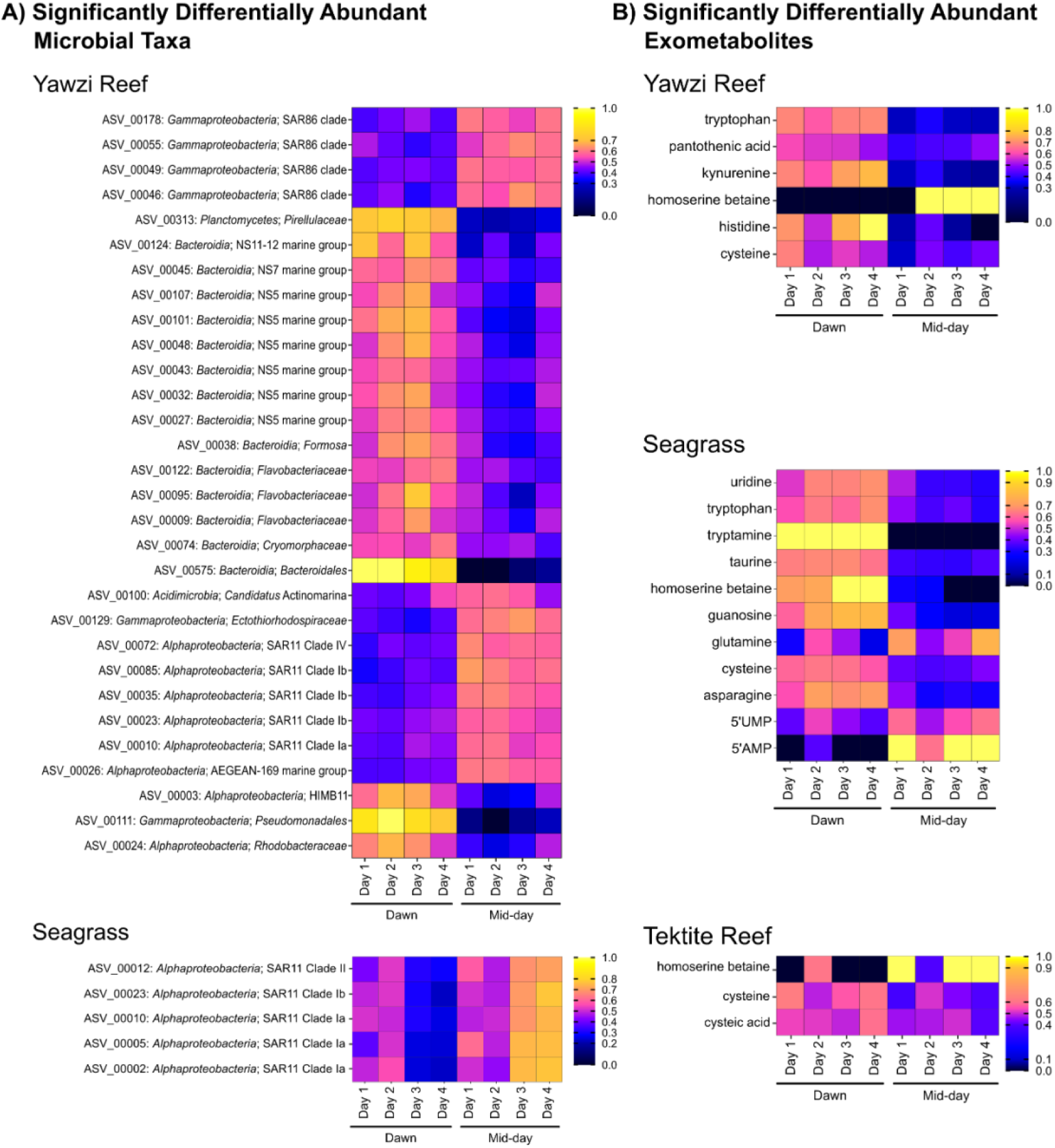
Temporally significant ASVs (A) and exometabolites (B) in Tektite Reef, Yawzi Reef, and Seagrass seawater. 33 unique ASVs were considered temporally significant based on concordance between corncob (5% FDR, p < 0.054) and DESeq2 (10% FDR, p < 0.050). 15 unique exometabolites were considered significant based on a Kruskal–Wallis test with FDR-adjusted p < 0.05. For each ASV and exometabolite, the median value across replicates at each time point and sampling day was calculated and then normalized to that day by dividing by that feature’s total daily abundance or concentration. Warmer colors indicate higher normalized abundance, and cooler colors indicate lower normalized abundance.

While exometabolome temporal dynamics were strongest in the seagrass and minimal at the two coral reefs, microbial variation was most apparent at Yawzi reef. The seagrass site showed limited microbial variation with time, with only five SAR11 ASVs, spanning clades Ia, Ib, and II, enriched at mid-day. At Tektite reef, no ASVs showed diel trends. At Yawzi reef, 30 ASVs varied significantly with time of day, including representatives of SAR11, SAR86, *Rhodobacteraceae*, *Flavobacteriaceae*, and other taxa. Only a small subset of SAR11 ASVs with significant temporal trends were observed at more than one site, highlighting a limited but recurring component of the microbial community that tracks diel changes.

### Hydrodynamics uniquely influence exometabolome and microbiome variability

Particle-tracking simulations were used to identify benthic particle origins and estimate the time evolution of the percentage of the water samples originating from the larger Lameshur Bay region (Figure 5). Particle source percentage in the seagrass, located within the bayhead, was highly stable with little to no exchange outside of the bay up to 15 hours before sampling each day (Figure S3). Conversely, the majority of particles at Tektite reef originated from outside Lameshur Bay, with less than 20% of particles originating from Lameshur Bay up to 24 hours prior to sampling (Figure S4), consistent with expectations given Tektite reef’s location at the southeast corner of the larger Lameshur Bay region. Unlike the seagrass site and Tektite reef, particle simulations at Yawzi reef showed significantly different water sources at dawn vs. mid-day, with particle origins outside Lameshur Bay observed as early as one hour before sampling (Figure S5). A higher percentage of Yawzi reef particles, on average, were from the larger Lameshur Bay region at dawn (43.4%) compared to mid-day (33.3%).

**Figure 5.**
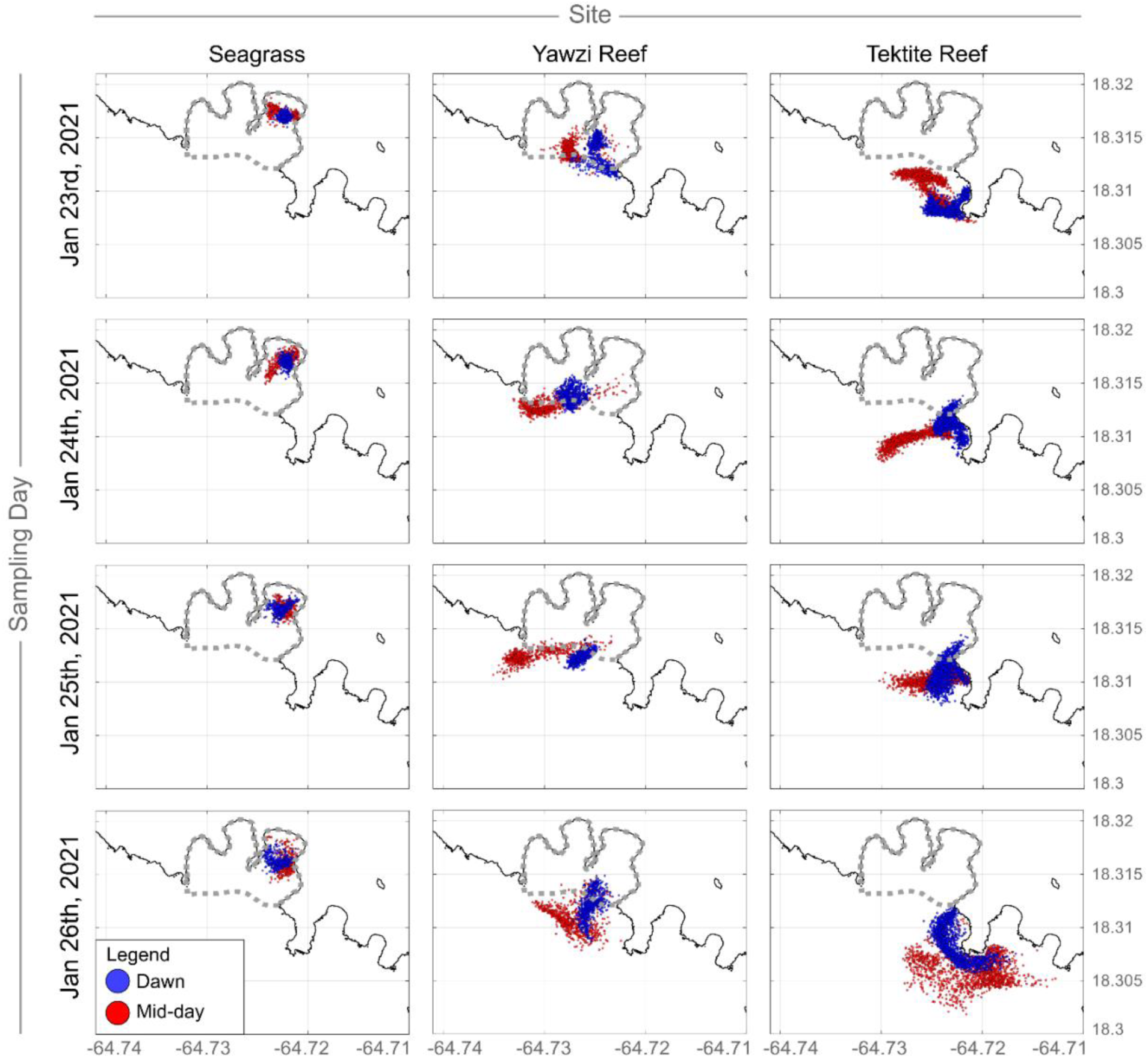
Hydrodynamics influence particle movement on diel timescales. Hydrodynamic and particle tracking models were used to trace the backward movement of benthic particles seven hours before exometabolome and microbiome sampling times. Particles were ‘released’ in the model between 5:00-7:00AM LST (dawn) and 12:00-2:00PM LST (mid-day) in 10-minute intervals. Particle-tracking simulations were carried out for each of the four days sampled in this study. Positions of the particles at 7 hours before the sampling times at dawn (blue) and mid-day (red) are shown by their respective latitude (y-axis) and longitude (x-axis) placements within the map. The boundary (dashed, grey line) represents the larger Lameshur Bay region.

Using a linear mixed model (Table S9), we found the microbiome was strongly influenced by the percent of particles in Lameshur Bay (regression coefficient [β] = 0.00022 ± 0.00008 SE, t = 2.75, R^2^ marginal = 0.342) in addition to a strong site influence (R^2^ conditional = 0.414). Additionally, beta-dispersion was positively associated with the percent of particles in both the linear regression (slope = +0.00024, adj. R² = 0.369, Figure S6b) and the linear mixed model (slope = +0.00022), indicating that particle origin contributed meaningfully to microbial variability even after accounting for site-level differences. In contrast, the percent of particles in Lameshur Bay did not influence the exometabolome dispersion (β = 0.00056 ± 0.00047 SE, t = 1.20, R^2^ marginal = 0.048), which was strongly influenced by site (R^2^ conditional = 0.893). A linear regression of beta-dispersion against the percent of particles in Lameshur Bay suggested a negative relationship for the exometabolome (slope = –0.0012, adj. R² = 0.535, Figure S6a). However, when accounting for site as a random effect in a linear mixed model, this relationship was no longer evident (slope = +0.00056), indicating that site-level environmental structure, rather than particles themselves, drove the observed exometabolome patterns.

### Relationships between exometabolites and microbes reflect ecosystem dynamics

To link changes in exometabolite concentrations and the abundance of microorganisms, site-level Spearman correlations were conducted using a Spearman’s rho cut-off of |0.5| and FDR q-value of 1%. A total of 1704 significant correlations were detected (Table S10). Specifically, 980, 442, and 373 significant correlations were detected for seagrass, Tektite, and Yawzi, respectively, between 42 exometabolites and 517 ASVs. Twenty-one of 517 ASVs had significant correlations with 10 or more exometabolites (Table S11), 3 of which were observed in more than one site, highlighting a high degree of specificity at the ASV level. Surprisingly, the taxonomy of these ASVs did not include typical photoautotrophic taxa. In contrast to the ASVs, individual exometabolites displayed greater redundancy, with pantothenic acid (vitamin B5) showing the highest number of correlations with ASVs (216; Figure 6b), while tryptamine had the fewest (9). Strong correlations (rho > |0.9|) were observed for 18 unique ASVs responsible for 19 ASV-metabolite relationships, 16 of which were found at Yawzi reef.

**Figure 6.**
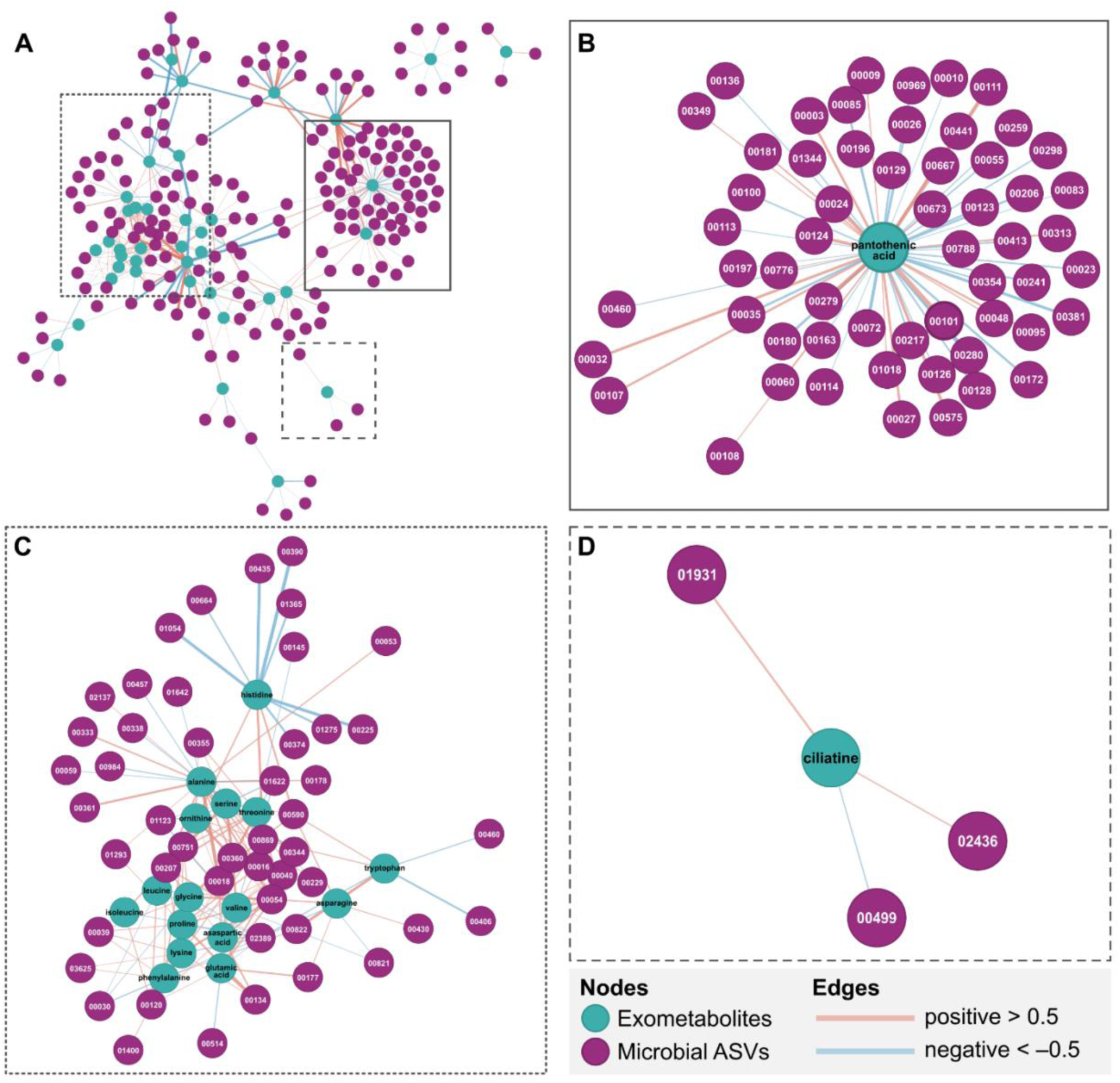
A) Complete correlation network showing all significant exometabolite-ASV correlations at Yawzi reef using Spearman correlations between exometabolite concentrations and ASV relative abundances. Correlations with a BH adjusted p-value < 0.05 or a significant p-value at the FDR Q = 1% were considered significant. Each node represents an exometabolite (green) or microbial ASV (purple). Edges, or connections between nodes, indicate the strength of the relationship, with thicker edges indicating a higher correlation coefficient (weight). Edge color represents the direction of the correlation, with positive correlations in red and negative correlations in blue. B) Zoomed in view of the pantothenic acid node and its connected microbial ASVs. C) Zoomed in view of a large sub-network composed of amino acid compounds and their connected microbial ASVs. D) Zoomed in view of the ciliatine node and its connected microbial ASVs.

## Discussion

By integrating quantitative exometabolomics, microbial community composition analysis, and hydrodynamic simulations over a short time-series, our results reflect patterns and potential drivers of exometabolome and microbial dynamics in Lameshur Bay. Specifically, patterns differed among sites, but all were affected by habitat type, time of day, and hydrodynamics, highlighting the complexity of interactions. Statistical analyses identified specific microorganism and exometabolite correlations, providing hypotheses for future studies to examine whether these linked indicators reflect coastal microbial processes.

Coral reef microbial communities were fairly homogeneous and characterized by taxa that are typically present in oligotrophic environments, including *Prochlorococcus, Synechococcus,* and SAR11. In contrast, the seagrass displayed a more diverse microbial community enriched with the phylum Bacteroidota, including *Flavobacteriaceae*, *Rhodobacteraceae*, *Cryomorphaceae*, and *Halieaceae*, which have previously been identified as core microbial taxa overlying seagrass [54]. Additionally, the sulfate-reducing bacteria *Desulfocapsaceae*, a microbe known to be associated with the seagrass rhizosphere [55–57], were enriched in the seagrass water. The consistent enrichment of these metabolically versatile groups suggests that seagrass meadows represent more organic-rich, dynamic habitats, supporting greater microbial diversity and functional potential than the surrounding reefs. While discernible differences between the coral reef and seagrass microbial communities were observed, only minor differences existed between the two reefs. Microbial community composition between the reefs differed more at dawn than at mid-day, with only two significant ASVs detected at mid-day - both elevated at Yawzi reef. However, the overall number of significant ASVs between the two reefs was markedly lower than between either reef and the seagrass, representing roughly a 5x and 50x decrease in significant ASVs at dawn and mid-day, respectively, relative to the seagrass. This consistency observed across adjacent reefs and over the span of multiple days suggests a largely uniform microbial community present among reefs within Lameshur Bay, consistent with prior findings [28]. Our results indicate that reef and seagrass microbial communities are primarily shaped by habitat, likely reflecting benthic composition and ecological function rather than short-term diel or day-to-day fluctuations.

Habitat type also served as the strongest driver of exometabolome composition. While average bulk total organic carbon measurements were comparable across habitats (Figure S2a), clear differences were observed in the concentration of individual amine- and alcohol-containing exometabolites, important labile components of DOM [8]. Seagrass exometabolomes were enriched in nucleotides, organosulfur compounds, and stress-protective osmolytes, whereas coral reef exometabolomes were dominated by proteinogenic amino acids. Corals have been shown to synthesize [58] and release dissolved free amino acids into the water column, via their mucus [59] or as dissolved exudates as a function of photosynthesis [60], at higher abundances than other benthic reef constituents [61]. The abundance of organosulfur compounds in the seagrass exometabolome could reflect metabolic strategies used by the seagrass plant to transform and release excess sulfide into organic forms, thereby allowing the plants to survive in sulfide-rich and anoxic sediments [62]. Experiments are needed to confirm this hypothesis. Overall, the majority of exometabolite concentrations were elevated in the seagrass habitat compared to the two coral reefs. We hypothesized that this concentration effect in seagrass reflected the more stable hydrodynamics of the meadow; however, linear mixed-model results revealed that exometabolome dispersion was more strongly driven by habitat than by hydrodynamics, likely reflecting the distinctive microbial and benthic composition of the two habitats.

By constructing correlation networks between exometabolites and microbial ASVs at Yawzi Reef (Figure 6a), we observed three general trends that warrant deeper investigation. First, many common metabolites, such as amino acids, are linked to numerous and diverse microorganisms, likely reflecting their central role in metabolism and nutrient cycling. For example, dissolved free amino acids constitute a major pool of labile dissolved organic nitrogen and are rapidly turned over by heterotrophic bacteria [63], consistent with their broad connectivity across microbial taxa (Figure 6c). Second, more selective associations emerged for exometabolite ciliatine, a phosphonate compound that contains a chemically stable carbon–phosphorus (C-P) bond and thus requires specialized metabolic pathways for degradation and assimilation. While phosphonates support a significant fraction of microbial P demand in the ocean, only two bacterioplankton species have been experimentally confirmed as phosphonate producers, and only a subset of marine microbes are capable of utilizing phosphonates, making these relationships more ecologically specialized (Figure 6d) [64, 65]. Finally, pantothenic acid, a vitamin precursor required for coenzyme A synthesis, is a highly conserved and ancient metabolite essential for many metabolic pathways, which likely underlies its extensive correlations with diverse microbial ASVs (Figure 6b) [66]. Collectively, these patterns present new hypotheses about metabolite exchange, nutrient use strategies, and functional interactions within coastal ecosystems.

Diel and tidal cycles shape the rhythm of coastal ecosystems, with fluctuations in light and water movement jointly regulating the activity, distribution, and interactions of marine microorganisms. In this study, the seagrass site showed highly stable hydrodynamics over a 24-hour time period, with little particle movement into or out of Lameshur Bay. This static environment resulted in strong correlations between exometabolites and microbial ASVs, as well as the most temporally significant exometabolites, including amino acids and their derivatives (e.g., tryptophan, glutamine, cysteine), nucleosides and nucleotides (e.g., uridine, guanosine, 5’-AMP, 5’-UMP), and osmoprotective compounds (e.g., taurine, homoserine betaine). In contrast, SAR11 (clades Ia, Ib, and II) was the only microbe to show significant temporal fluctuations between dawn and mid-day. SAR11 was consistently elevated at mid-day in the seagrass, consistent with prior observations where SAR11 abundances were increased in the evening compared to the morning or afternoon [67]. Among the two coral reefs, homogenous exometabolome and microbiome profiles were observed at each distinct sampling time, but with varying temporal dynamics and significantly fewer correlations compared to the seagrass environment. Specifically, more pronounced temporal changes were observed at Yawzi compared to Tektite. Yawzi also exhibited the unique trend where exometabolite profiles were significantly more diverse at mid-day, during the photosynthetically active diel period, while the microbial community, conversely, was less diverse. We hypothesized that hydrodynamics might be strongly influencing the microbial communities at Yawzi reef through mixing or flushing. Our hydrodynamic model demonstrated low mid-day presence of coastal bay water at Yawzi reef due to a large influx of offshore water, which likely caused exometabolite and microbial community variability and a more homogenous influx of offshore/oligotrophic microbes at mid-day. This influx of offshore water and more dynamic environment at Yawzi may also explain the decrease in significant exometabolite-microbe correlations compared to Tektite, where a more stable hydrodynamic environment may improve detection of these relationships.

Our results revealed that habitat type, time of day, and hydrodynamics all strongly influence the microbial communities and exometabolites present within coral reefs and seagrass meadows. While current monitoring strategies rely heavily on visual observations and macro - organismal metrics, our findings highlight the capabilities of ‘omics measurements to detect subtle shifts in the daily rhythms of exometabolite–microbial interactions, and that hydrodynamical modeling is useful to further explain the trends. Integrating exometabolites, microorganisms, and hydrodynamics offers a path toward more proactive and informative monitoring frameworks and should be considered in the design of restoration strategies.

## Associated Content

### Data Availability

All MS data are available at MetaboLights under accession number MTBLS9008 (https://www.ebi.ac.uk/metabolights/MTBLS9008) and the sequencing data are available at NCBI SRA under accession PRJNA1380722. A step-by-step BC derivatization protocol is publicly available on protocols.io (dx.doi.org/10.17504/protocols.io.biukkeuw). All MATLAB® scripts used for processing the Skyline outputs are available on GitHub (https://github.com/KujawinskiLaboratory/SkyMat).

### Author Contributions

- Study design – Amy Apprill, Elizabeth Kujawinski, Laura Weber, W. Gordon Zhang
- Permit - Amy Apprill
- Sample collection - Laura Weber, Cynthia Becker, Amy Apprill
- Sample processing - Laura Weber, Cynthia Becker, Gretchen Swarr, Mallory Kastner, Aushaun Brown
- Instrumental analysis - Melissa C. Kido Soule
- Data processing - Brianna Garcia, Sharon Grim, Yan Jia
- Data analysis - Brianna Garcia, Sharon Grim
- Writing - Brianna Garcia, Sharon Grim, Amy Apprill
- Editing - All authors
- Funding - Amy Apprill, Elizabeth Kujawinski

### Funding

This work was supported by National Science Foundation OCE awards 1736288, 2414888, and 2307424 and a NOAA OAR Cooperative Institutes award to AA and EK (#NA19OAR4320074).

## Acknowledgments

We thank Nadège Aoki and Justin Ossolinski for assisting with field sampling and processing, Krista Longnecker for assisting with TOC/TN sample processing and analysis, the Roy J. Carver Biotechnology Center of the University of Illinois at Urbana-Champaign for DNA sequencing, and Erin McParland for helpful discussions regarding experimental design. The following reagent was obtained through BEI Resources, NIAID, NIH as part of the Human Microbiome Project: Genomic DNA from Microbial Mock Community B (Even, Low Concentration), v5.1L, for 16S rRNA Gene Sequencing, HM-782D. Samples were collected using National Park Service permit VIIS-2021-SCI-0002. This work was supported by National Science Foundation OCE awards 1736288, 2414888, and 2307424, NOAA OAR Cooperative Institutes award to AA and EK (#NA19OAR4320074) and WHOI’s Reef Solutions Initiative.

