## Supplemental Text for "Habitat and Hydrodynamics Influence Coral Reef and Seagrass Microbial and Exometabolite Dynamics"

**Author emails and ORCIDs:**
Aushaun Brown -

**Affiliations:**
[1]Department of Marine Chemistry and Geochemistry, Woods Hole Oceanographic Institution, Woods Hole, Massachusetts, United States;
[2]Department of Applied Ocean Physics & Engineering, Woods Hole Oceanographic Institution, Woods Hole, Massachusetts, United States;
[3]The MIT-WHOI Joint Program in Oceanography/Applied Ocean Science and Engineering, Cambridge and Woods Hole, Massachusetts, United States;
[4]Department of Biology, Ithaca College, Ithaca, New York, United States;
[5]University of the Virgin Islands, 2 John Brewers Bay, St. Thomas, U.S Virgin Islands

**Supplemental Methods:**

Materials:

Hydrochloric acid (HCl), acetone, methanol (MeOH), acetonitrile (MeCN), benzoyl chloride (BC; 99%, ACROS Organics), and sodium hydroxide (NaOH) were purchased from Fisher Scientific (Optima). Phosphoric acid (85%, ACS reagent grade), $^{13}C_6$-ring-BC (99% atom $^{13}C$), and internal standards were purchased from Sigma-Aldrich. Stable isotopically labeled - internal standards matched to each targeted compound (SIL-IS) were prepared with $^{13}C_6$-ring-BC. Deionized water was obtained from a Milli-Q system (Millipore; resistivity 18.2 MΩ at 24 °C, TOC < 1 μM). Samples and reagents were stored in acid-washed, combusted (at least 4 h at 450°C) glassware. BC and all solvents were transferred using combusted glass Pasteur pipettes. Primary stocks and mixes were stored at −20°C. The working reagent (5% BC in acetone) was prepared fresh daily.

Total Organic Carbon and Microbial Abundances:

From each Niskin water collection, 40 mL of unfiltered benthic seawater was processed and analyzed for total organic carbon (TOC) and total nitrogen (TN) concentrations, and 1.4 mL was collected, processed, and analyzed for the enumeration of *Prochlorococcus*, *Synechococcus*, picoeukaryotic cells, and unpigmented cells (generally heterotrophic bacteria and archaea) following methods described elsewhere [1].

Targeted metabolomics UHPLC-MS/MS data collection:

Reverse phase (RP) chromatography was performed according to Widner et al [2]. Briefly, using an ultra-high performance liquid chromatography system (UHPLC; Vanquish, Thermo Scientific, Waltham, MA, USA), separation was achieved on a Waters Acquity HSS T3 column (100Å, 1.8 μm particle size, 2.1 mm X 100 mm) equipped with an Acquity HSS T3 VanGuard pre-column (100Å, 1.8 μm particle size, 2.1 mm X 5 mm) coupled to a heated electrospray ionization source (H-ESI) and an Orbitrap mass spectrometer (Orbitrap Fusion Lumos, Thermo Scientific).

Chromatography gradient, mass spectrometer settings, and quality assurance and control (QA/QC) are described in detail elsewhere [1].

Metabolomics data processing:

Thermo .raw files were processed using Skyline (v. 21.2.0.568) [3, 4]. Peaks corresponding to detected metabolites were integrated based on accurate mass (± 10 ppm), retention time, and MS/MS fragment ion confirmation. Calibration curves for each compound were constructed based on the amount of metabolite standard added (ng-added) versus the integrated light-to-heavy peak area ratios. Exported quantification tables from Skyline containing peak areas for each metabolite were imported into MATLAB® (v. R2022a). An in-house MATLAB® script (considerSkyline.m) was used to generate linear regression calibration curves, quantify each metabolite, calculate limits of detection (LOD) and quantification (LOQ), and merge positive and negative polarity mode data. All MATLAB® scripts used for processing the Skyline output are available at https://github.com/KujawinskiLaboratory/SkyMat. The resulting merged quantification table was further filtered based on the QA/QC metrics described below. The resulting quantification table containing nanomolar (nM) concentrations was $\log_2(x+1)$ transformed for downstream statistical analyses to handle the dynamic range of metabolite concentrations appropriately and avoid negative transformed values due to imputation of values less than the LOD.

Quality Assurance and Quality Control (QA/QC):

Thorough QA/QC was conducted as described by Garcia et al. 2024 [1] to ensure the reliability of the metabolites reported herein, with 45 metabolites retained post-QA/QC. Sample outliers were determined through a combination of calculations which included (1) the Euclidean distance between each sample within a site at a given sampling time, (2) the 95% confidence interval (CI) for each site/sampling time non-metric multidimensional scaling (NMDS) plot, and (3) Bray-Curtis dissimilarity distance from the site average within a given sampling time. Sample outliers were removed conservatively, as biological variation across replicates was anticipated. Sample #73 was removed as an outlier as it deviated outside the 1.5 times the interquartile range +/- the upper (75th) quartile and fell outside the 95% confidence interval of the seagrass mid-day sampling group and clustered on top of the coral reef sample groups.

16S rRNA gene data processing:

Amplicon libraries were analyzed using the DADA2 (v. 1.26.0) package in R. Read pairs were assessed for quality, trimmed of primers, and used to estimate sequencing error rates (filterAndTrim: truncLen = c(210, 160), maxN = 0, matchIDs = TRUE, multithread = 12, maxEE = c(2,2), truncQ = 2, rm.phix = TRUE, compress = TRUE). Read pairs were merged to cover the length of the 16S rRNA gene V4 region (mergePairs: minOverlap = 12), and the resulting amplicon sequence variants (ASVs) were checked for chimeras (removeBimeraDenovo: method = "consensus", multithread = nthreads). Taxonomy was assigned to ASVs using the Silva v.138.1 database (https://zenodo.org/records/4587955) (assignTaxonomy: minBoot = 50, tryRC = TRUE, multithread = 12). R package decontam (v. 1.18.0) was used to identify putative contaminant sequences using negative control samples from DNA extraction and PCR amplifications (isContaminant: method = "minimum", threshold = 0.1). We examined the distribution of putative contaminants at both the species level and ASV level, to account for endemism and biological variability between sites. This method identified 339 potential contaminant ASVs representing 104 different species. After removing sequences assigned to potential contaminants (n = 7,617) and of eukaryotic origin (chloroplasts, mitochondria, and eukaryotes: n = 171,607), the dataset of 90 samples retained 7,854,297 sequences representing 12,350 ASVs. Due to the nuances of taxonomic databases, for several ASVs taxonomic assignment was less specific beyond the class level. In order to include such ASVs that were also abundant in the study, the last known taxonomic lineage was propagated to the genus level. For example, of the 498 ASVs assigned to the Pseudomonadales family (Gammaproteobacteria) 269 had taxonomic assignments to the genus level. This excluded 49 abundant ASVs assigned at the family level to the SAR86 clade. Thus, to examine taxonomic diversity at ecologically relevant scales, the family-level SAR86 clade was also included.

<u>Statistical Analysis:</u>

*Metabolomics statistical analyses*

Statistical analyses were carried out in R Studio (2024.09.1+394) running R (4.3.2). Data was tested for normality visually via histogram plots and mathematically using the Shapiro–Wilk test (stats::shapiro.test). Most exometabolites were not normally distributed (p-value < 0.05) and thus non-parametric tests were used for all downstream analyses. Bray Curtis dissimilarity between metabolomic samples was calculated and visualized by non-metric multidimensional scaling (NMDS) using the 'vegan' [5] and 'ggplot2' [6] packages. To accommodate the wide range of inter-site diversity, we compared within-site NMDS visualizations from a distance matrix using all samples, as well as site-specific distance matrices. A Permutational Multivariate Analysis of Variance (PERMANOVA) using Bray Curtis distance matrices was carried out on log2(x+1) exometabolite concentrations and environmental categorical variables using the adonis2 function in the ‘vegan’ package. A distance-based redundancy analysis (dbRDA) was used to test categorical and continuous environmental and biological factors against the metabolomics data using Bray-Curtis distances calculated by vegan::vegdist and the vegan::capscale functions. A Kruskal-Wallis test (stats::kruskal.test) was used to test for significant exometabolites between dawn and mid-day samples within sampling sites. A Kruskal-Wallis test (stats::kruskal.test) followed by Pairwise Wilcoxon Rank Sum tests (stats::pairwise.wilcox.test) were conducted to determine significant exometabolites across sites within a sampling time. P-values were adjusted to account for multiple comparisons using the Benjamini-Hochberg correction using an implementation of stats::p.adjust(method = “fdr”), and exometabolites were considered significant with a false discovery rate (FDR) adjusted p-value < 0.05.

*Microbial statistical analyses*

Amplicon sequence data were analyzed in RStudio (v 2024.09.0) running R (v4.2.2) with the following packages: tidyverse (v 2.0.0), phyloseq (v 1.42.0), corncob, DESeq2, vegan (v 2.6.1), ashr (v. 2.2.63), boot (v. 1.3.31), spiec.easi (v 1.1.3), and NetCoMi (v 1.1.0). Alpha-diversity estimates (vegan::rarecurve) (vegan v 2.6.1) for samples within each site used per-sample raw sequence counts rarefied to the lowest sequence depth observed within each site. Morisita-Horn dissimilarity distances (vegan::metaMDS) were calculated between samples' relative abundances of the amplicon sequence variants (ASVs), which performed better than other metrics (e.g. Bray-Curtis). As with the exometabolomes, NMDS visualizations were constructed from a distance matrix using all samples, as well as site-specific distance matrices to accommodate the wide range of inter-site diversity. R packages corncob (v. 0.4.1) and DESeq2 (v. 1.38.3) were used to identify significantly differentially abundant ASVs between pairs of sites at the same sampling time, and between sampling times within the same site (total of 9 contrasts). Each package handles data and statistical testing differently; corncob uses one of three tests (Wald, log-likelihood ratio, or Rao) to examine differential abundance and variability in abundance of ASVs between groups of samples. By default, corncob filters for significant taxa using an implementation of stats::p.adjust(method = "fdr"), which assumes the observations are independent. A more conservative approach was used to filter the results, by modeling the local false discovery rates (LFDR) (ashr::qval.from.lfdr) from the computed p-values, and calculating the 1%, 5%, and 10% LFDR quantiles. After examining the p-values distributed in those quantiles, the corresponding adjusted p-values, modeled estimates and coefficients for all observed taxa, a 5% ($\alpha = 0.05$) experiment-wise error rate was determined to be most effective at culling putative false positives that arose from multiple tests of non-independent observations. This post-test filtering step retained 708 observations of 207 unique ASVs, with a maximum corresponding p-value of $p < 0.0539$). DESeq2 normalizes ASV relative abundance and sample abundance by sampling depth and number of samples, sets the per-ASV expected abundance (null hypothesis) as the mean abundance across the input samples, and can implement a variety of metrics for post-test corrections beyond constraining the false discovery rate. Similar to post-test handling of corncob results, raw p-values from DESeq2::results(alpha = 0.05) were used to model 1%, 5%, and 10% LFDR quantiles. An experiment-wise false discovery rate of 10% ($\alpha = 0.10$) was used to filter the corresponding adjusted p-values and shrunken log2(fold change) estimates from DESeq2::lfcShrink(type = "ashr"), yielding 1306 observations of 370 unique ASVs with a maximum corresponding p-value of $p < 0.0497$. Results from both methods were compared; 189 distinct ASVs unique to DESeq2 and 26 distinct ASVs unique to corncob were identified as significantly differentially abundant with respect to method. Across both methods, 181 unique ASVs were flagged to be significantly differentially abundant in paired sample groups, which we retained for further analyses.

*Concurrent exometabolome and microbiome diversity analyses:*

For exometabolomes and microbiomes, a permutation of analysis of variance (PERMANOVA) via vegan::adonis2 was calculated to assess the impact of site, sampling time, and sampling day on exometabolite and microbial population diversity. As with the NMDS, the input pairwise dissimilarity matrices to PERMANOVA were constructed from Bray-Curtis distances on log2(x+1) transformed exometabolite profiles, and Morisita-Horn distances on ASV relative abundances. Estimates of variation were calculated from individual terms (site, time, and day), as well as interactions of terms. Interaction terms “sampling time x site” and “sampling time x sampling day x site” yielded negative estimates in the original model. As recommended, they were pooled one at a time with the term with the lowest estimate of variation, “sampling time x sampling day”, and the model was re-run until no negative estimates were generated [7]. The final model included the interaction terms “sampling day x site” and a pooled term (“sampling time x sampling day”, “sampling time x site”, and “sampling time x sampling day x site”), in addition to the original source variables. To estimate the 5% FDR threshold for significance, a custom permutation model using the boot::boot function generated a modeled distribution of p-values calculated from 9999 permutations of randomly subsampled data.

The impact of site, sampling time, and sampling day on microbial and exometabolite community diversity was assessed using bootstrapped t-tests (via stats::t.test) and Kruskal-Wallis rank sum tests (via stats::kruskal.wallis). For each comparison, the mean difference in beta diversity, here reported as $\Delta\beta$, was calculated from groups of sample beta diversity values. The distribution of beta diversities was assessed using vegan::betadisper(type = "median", sqrt.dist = FALSE), which calculates the dispersion or distance between each sample's coordinates in NMDS and a group representative centroid, in this study using the spatial median. The median of beta dispersion values for each group is referred to here as the median $\sigma_{\beta}^{2}$. Using beta diversity values quantifies similarity between groups of samples, whereas beta dispersion comparisons convey if the degree of variability within a specific group of samples is statistically different from another group.

A linear model (stats::lm) and linear mixed model (via lme4::lmer) were used to determine the influence of the particle source on the exometabolome and microbiome variability using average beta-dispersion distances previously calculated for each site (Table S9). Given that the dawn and mid-day sampling times were separated by ~7h, the average percent of particles within the greater Lameshur Bay region was calculated at each site seven hours before sampling across the four sampling days (Figure S3-S5). Linear regressions were constructed based on the linear model outputs (Figure S6). Site was included as a random effect in the linear mixed model.

*Exometabolite-microbe correlations:*

Site-specific spearman correlations were conducted between log2(x+1) transformed exometabolite concentrations and ASV abundances. Because some microbial samples had technical replicates in the exometabolite samples (one filter corresponding to multiple filtrate samples), technical replicate exometabolome samples were averaged to ensure a one-to-one relationship between the exometabolome and microbiome data. Due to the increased granularity of subsetting the site-specific datasets for correlation analyses, the impact of data granularity was assessed on adjusted p-values (stats::p.adjust) and FDR q-values (ashr::qval.from.lfdr) (Table S12), demonstrating that with increased granularity of the site-specific datasets that BH-adjusted p-values $< 0.05$ corresponded to a maximum raw p-value $< 0.0003$. For the explorative nature of this analysis, this level of stringency was deemed unnecessary. Thus, a Spearman's rho ($\rho$) of $|0.5|$ and the p-value corresponding to a 1% FDR were selected as the criteria for significant correlations.

Hydrodynamic modeling:

The St. John hydrodynamic model is based on the widely used Regional Ocean Modeling System (ROMS) and has been used to simulate fine-scale circulation in the St. John area, including the numerous bays, with a horizontal grid resolution of 50 m [8]. The model resolves the fine-scale coastal bathymetry, including the shallow shelf, around St. Thomas, St. John, and the British Virgin Islands. To simulate the time-evolving, three-dimensional flows around the islands and in the bays, the model captures vertical water mixing as well as bottom friction. The model is forced on the lateral open boundaries by calibrated realistic fields of tides, temperature, salinity, and velocity from large-scale ocean models and at the surface by hourly meteorological conditions from a global atmospheric model. Results of the model hindcast simulation in 2016-2022 were validated against historical observations on the St. John coastal region.

### Supplemental Results:

Sampling Day Significance

Samples were collected for exometabolome and microbial sequencing analysis over four consecutive days: January 23rd – January 26th, 2021. The PERMANOVA models suggested that a minor, but significant contribution to variability in the exometabolome (11.7%) and microbiome (8.3%) was site-specific sampling day (day x site, $p < 0.05$). More broadly, sampling day was found to significantly contribute to the microbiome (14.7% of variability, p-value < 0.001), yet not the exometabolome (p-value = 0.246). Individual metabolites were tested for significance across sampling days and revealed no significant metabolites (KW adj. P-value < 0.05). Sampling day was insignificant in the metabolomics dataset and had the lowest percent contribution (14.7%) of the investigated variables to the microbiome data, thus allowing for samples to be grouped across days based on site and/or sampling time allowing for robust statistics.

Environmental factors

A redundancy analysis (RDA) investigated the influence of various environmental and biological variables on the metabolome composition. Variables tested included total organic carbon (TOC), total nitrogen (TN), and abundances of *Prochlorococcus*, *Synechococcus*, picoeukaryotes, and heterotrophic cells. All measurements were log2(x+1) transformed and tested for colinearity before running the RDA. A strong positive correlation was found between *Prochlorococcus* and *Synechococcus* (0.87), therefore *Synechococcus* measurements were excluded from the RDA to limit variable redundancy. The significance of the individual terms in the RDA model was evaluated using an ANOVA and found no significant variables (p-value < 0.05) excluding site and sampling time, consistent with previous findings in the PERMANOVA.

**Supplemental Figures:**

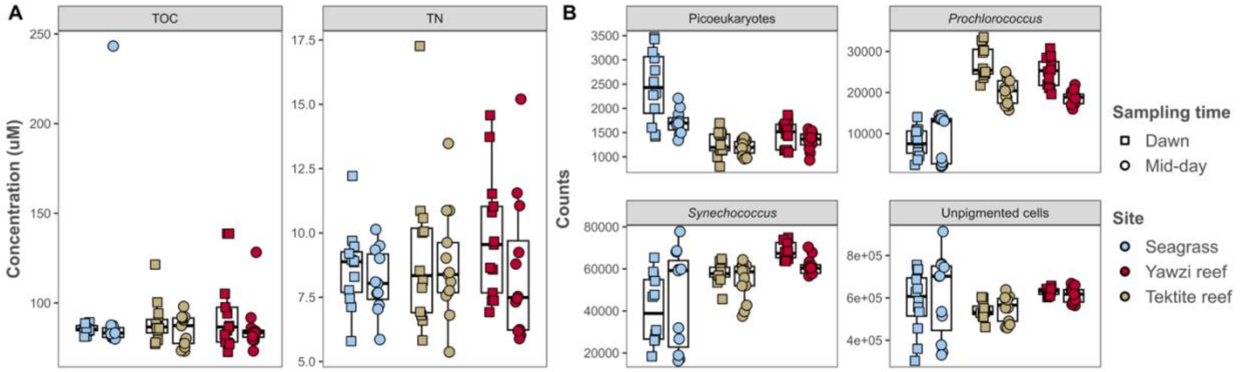


**Supplemental Figure 1.** Boxplots represented for each sampling site at dawn (squares) and mid-day (circles) over the four consecutive sampling days for (A) bulk water chemistry concentrations in micromolar (uM) of total organic carbon (TOC) and total nitrogen (TN), and (B) flow cytometry counts of picoeukaryotes, *Prochlorococcus*, *Synechococcus*, and unpigmented cells (heterotrophic bacteria and archaea).

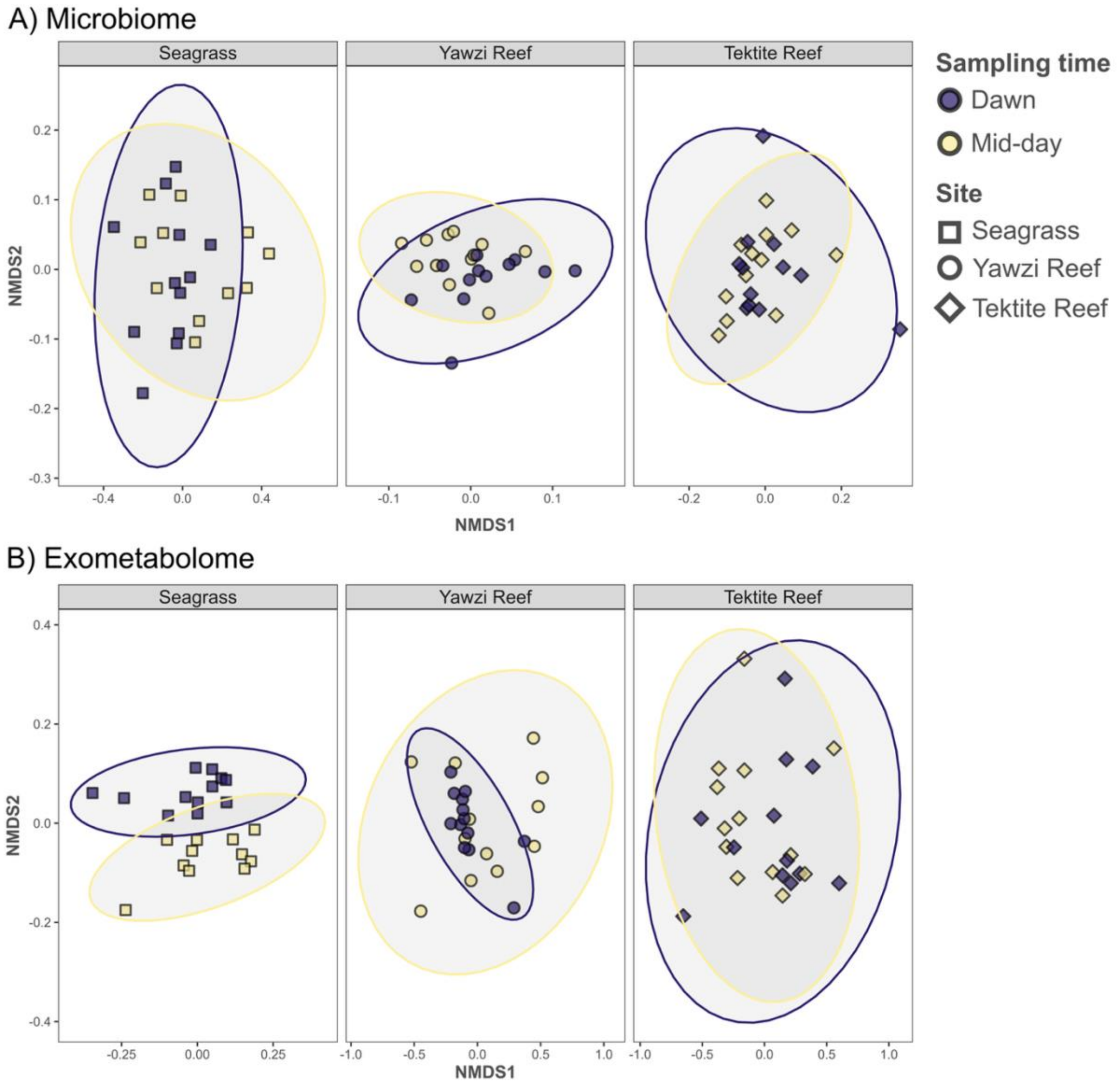


**Supplemental Figure 2. Site-specific NMDS ordination plots** of the (A) microbiome and (B) exometabolome are shown using Morisita-Horn and Bray-Curtis dissimilarities, respectively. Confidence intervals (95%) are shown as ellipses for dawn (blue) and mid-day (yellow) sampling times. Sites are separated into their own individual facets and data points are represented by unique shapes per site.

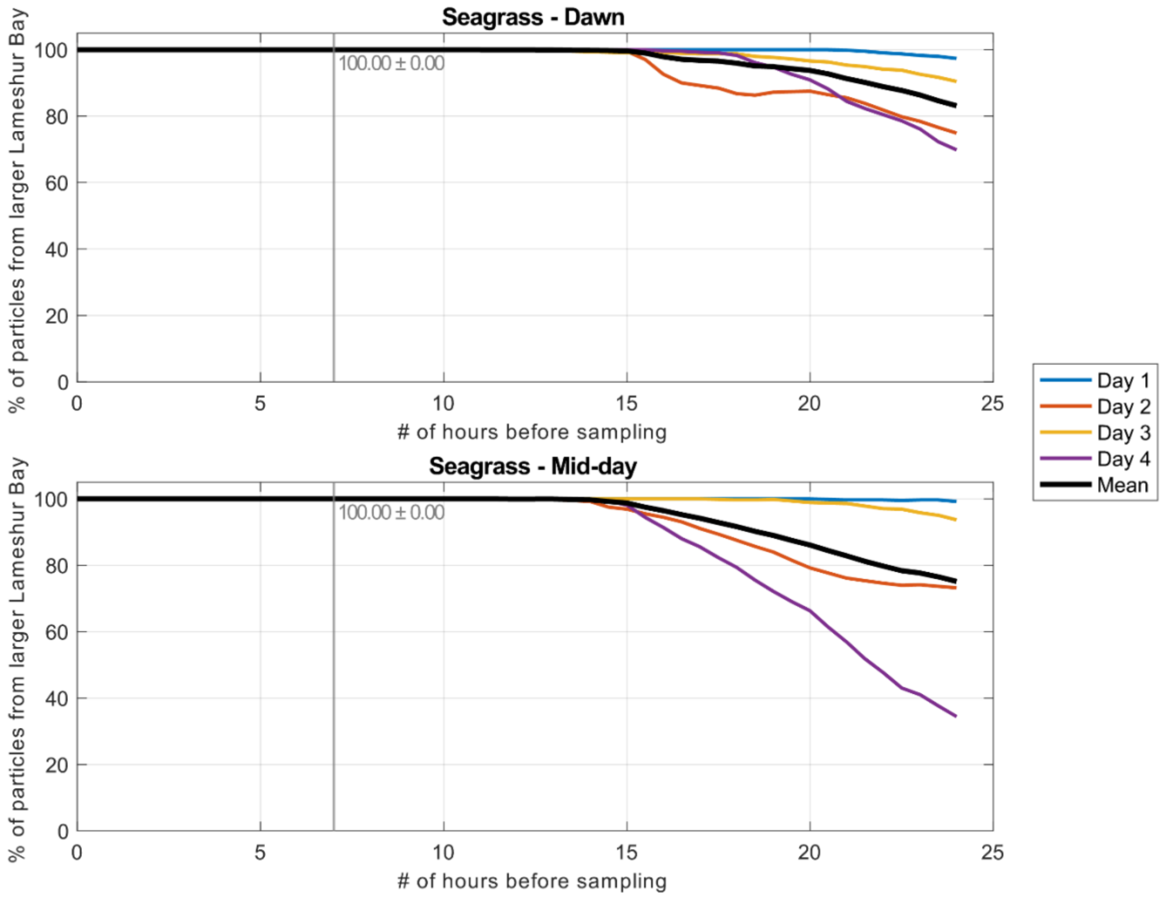


**Supplemental Figure 3. Sources of water samples in the Seagrass.** Hydrodynamic and particle-tracking models were used to identify the origin of the water samples. Here we show the percentage of particles from the larger coastal region of Lameshur Bay (y-axis) looking backwards in time where the number of hours prior to sampling is depicted on the x-axis. Particle-tracking simulations were carried out for each of the four sampling days, with the average shown in solid black. The average percentage of particles from the larger Lameshur Bay region at seven hours prior to sampling (the approximate number of hours between our sampling times) is shown as a vertical grey line with the average and standard deviation values of that timepoint shown on the plot. Separate simulations were constructed for the dawn and afternoon sampling periods.

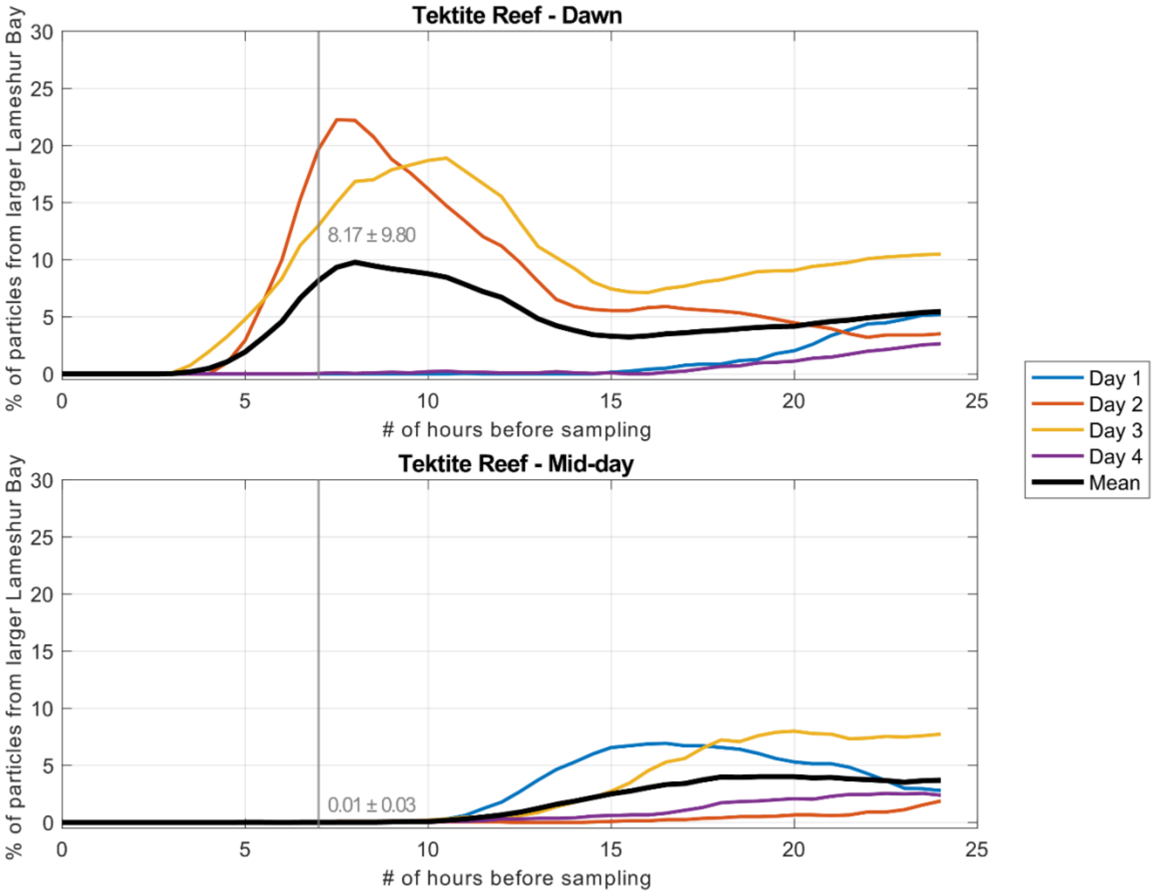


**Supplemental Figure 4. Source of water samples at Tektite Reef.** Hydrodynamic and particle-tracking models were used to identify the origin of the water samples. Here we show the percentage of particles from the larger coastal region of Lameshur Bay (y-axis) looking backwards in time where the number of hours prior to sampling is depicted on the x-axis. Particle-tracking simulations were carried out for each of the four sampling days, with the average shown in solid black. The average percentage of particles from the larger Lameshur Bay region at seven hours prior to sampling (the approximate number of hours between our sampling times) is shown as a vertical grey line with the average and standard deviation values of that timepoint shown on the plot. Separate simulations were constructed for the dawn and afternoon sampling periods.

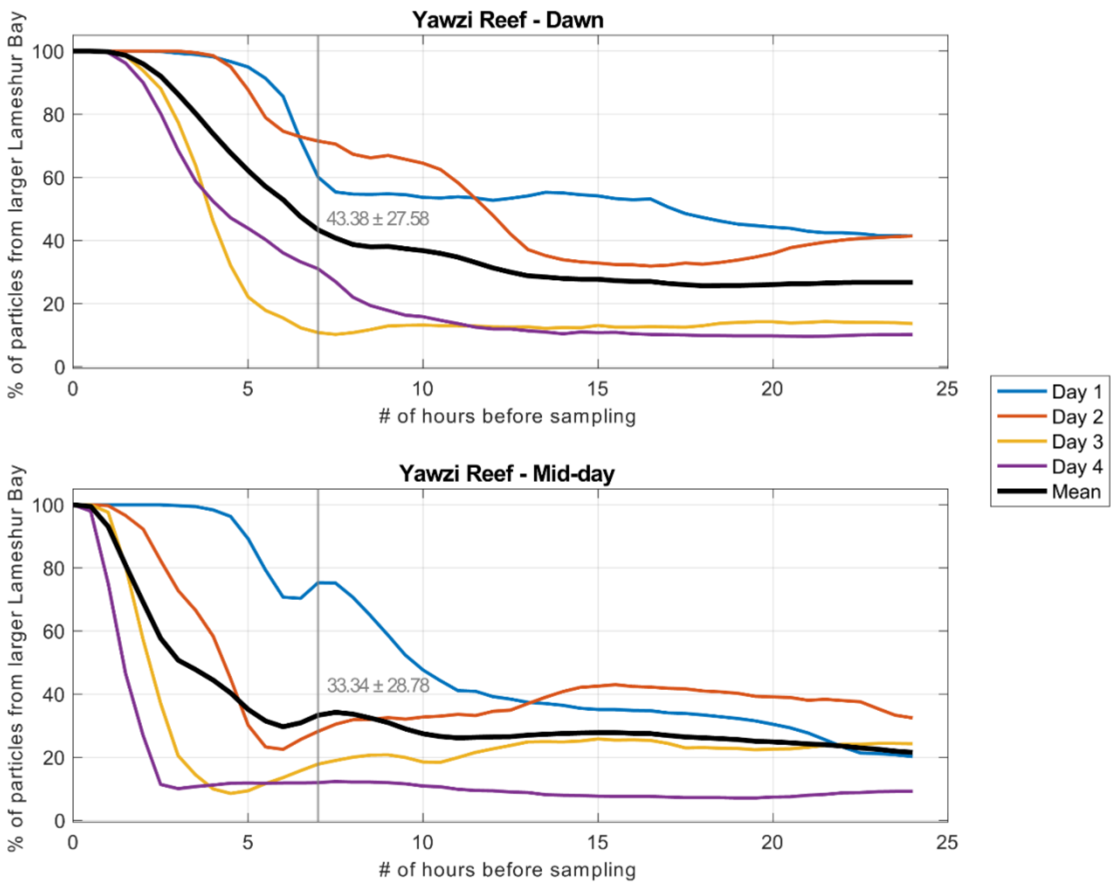


**Supplemental Figure 5. Source of water samples at Yawzi Reef.** Hydrodynamic and particle-tracking models were used to identify the origin of the water samples. Here we show the percentage of particles from the larger coastal region of Lameshur Bay (y-axis) looking backwards in time where the number of hours prior to sampling is depicted on the x-axis. Particle-tracking simulations were carried out for each of the four sampling days, with the average shown in solid black. The average percentage of particles from the larger Lameshur Bay region at seven hours prior to sampling (the approximate number of hours between our sampling times) is shown as a vertical grey line with the average and standard deviation values of that timepoint shown on the plot. Separate simulations were constructed for the dawn and afternoon sampling periods.

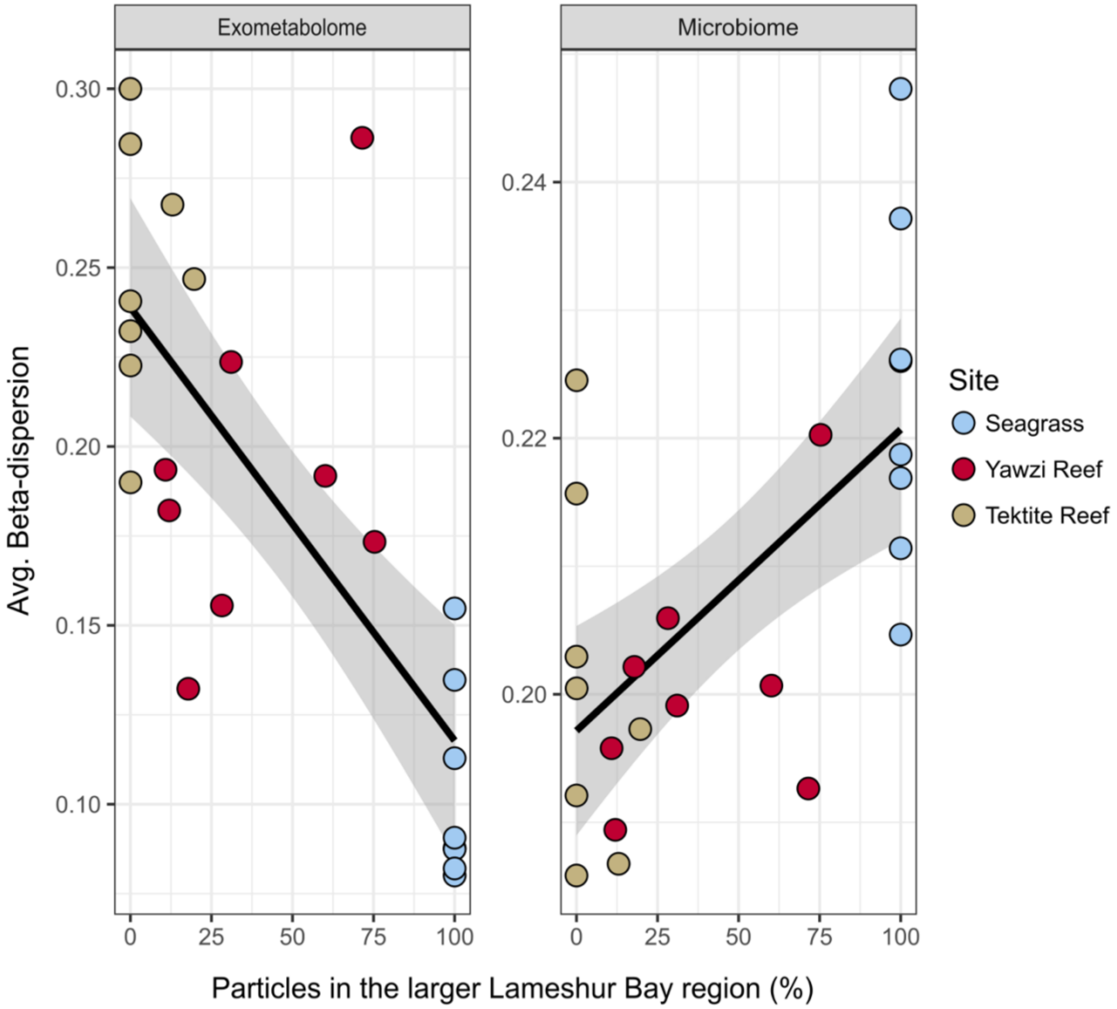


**Supplemental Figure 6.** Linear model of the average percent of particles from the larger Lameshur Bay region 7-hours prior to water sampling based on the hydrodynamic and particle-tracking models compared against the calculated average beta-dispersion at each sampling site for the exometabolome and microbiome.
